# The Virtual Biotech: A Multi-Agent AI Framework for Therapeutic Discovery and Development

**DOI:** 10.64898/2026.02.23.707551

**Authors:** Harrison G. Zhang, Peter Eckmann, Jiacheng Miao, Andrew B. Mahon, James Zou

## Abstract

Drug discovery and development requires integrating diverse evidence across biological scales and data modalities. However, relevant data, tools, and expertise remain fragmented across teams and organizations, making integration difficult. To address these challenges, we introduce the Virtual Biotech, a coordinated team of AI agents that mirrors the structure of human therapeutic research organizations to support end-to-end computational discovery. The Virtual Biotech is led by a Chief Scientific Officer agent that receives scientific queries, delegates them to domain-specialized scientist agents, and integrates their outputs through data-driven reasoning. Scientist agents leverage complementary tools and knowledge sources spanning statistical genetics, functional genomics, pathways and interactions, chemoinformatics, disease biology, and clinical data. We showcase the Virtual Biotech across three translational applications. First, the agents autonomously annotated and analyzed outcomes from 55,984 clinical trials to identify genomic features of drug targets associated with trial success. More than 37,000 clinical-trialist agents curated structured trial outcomes and linked targets to multi-omic annotations, including cell-type-specific features derived by the agents from single-cell RNA-sequencing atlases. The agents discovered that drugs targeting cell-type-specific genes were 40% more likely to progress from Phase I to Phase II and 48% more likely to reach market (Phase IV), while exhibiting 32% lower adverse event rates. Second, the Virtual Biotech evaluated B7-H3 as a lung cancer target, integrating statistical genetics, single-cell, spatial, and clinicogenomic evidence to propose an antibody–drug conjugate strategy while identifying key liabilities and differentiation opportunities. Third, the platform analyzed a terminated ulcerative colitis trial targeting OSMR*β* to infer potential failure mechanisms and proposed biomarker-guided enrollment strategies to address precision-medicine gaps. Together, these results illustrate how the Virtual Biotech can enable more transparent, efficient, and comprehensive multi-scale therapeutic analyses, helping to accelerate early-stage drug discovery workflows while keeping human scientists in the loop.

## 1 Introduction

Drug discovery is one of the most consequential and challenging endeavors in biomedicine. Bringing a single therapeutic from concept to clinic often requires more than a decade of development and costs on the order of billions of dollars [1–3]. Despite these investments, approximately 90% of drug candidates entering Phase I trials never reach approval, with failures most often driven by insufficient efficacy and unexpected safety liabilities [4–7]. To reduce this attrition, modern drug discovery increasingly draws on evidence from diverse sources to validate targets, including but not limited to human genetics, functional genomics, single-cell and spatial profiling, structural biology, medicinal chemistry, and clinical medicine [8].

However, there are a number of organizational and scientific bottlenecks that make use of this evidence challenging in practice. First, the expertise to collect and analyze different sources of evidence is fragmented across disciplines, formats, and biological scales. This fragmentation leads to siloed workflows across teams and organizations, missing opportunities for nuanced cross-referenced findings [3, 9, 10]. Second, making decisions based on disparate and sometimes conflicting sources of evidence is difficult, even more so when non-data driven factors like organizational dynamics, differing prior beliefs, or employee incentives can influence decisions [3, 9]. Third, findings can be difficult to audit as fragmented teams each document their discovery process differently, making it hard to understand why a decision was made [3, 11, 12]. Lastly, the recent explosion in the scale of biological data renders manual analysis and integration of multiple modalities difficult and time-consuming [3, 9]. Thus, despite advances in data-driven approaches, early stage drug development remains slow, resource intensive, and prone to failure [1–7].

Artificial intelligence (AI) agents have been proposed to accelerate and scale scientific tasks that involve extensive data analysis. Agents are large language model (LLM) systems equipped with external resources including tools and databases, giving them the ability to autonomously carry out complex multi-step tasks and reasoning. By giving an agent a clearly specified role and equipping it with tools tailored to the task (for instance, bioinformatics analysis software), the agent can be tailored to address a specific category of problems. Recent work also explores multiagent systems, where multiple specialized agents collaborate to solve complex problems [13].

Here, we introduce the Virtual Biotech, a multi-agent AI research platform designed to comprehensively and efficiently generate evidence to answer scientific queries in the early drug discovery pipeline. At the core is a virtual Chief Scientific Officer (CSO) orchestrator, which breaks down queries into manageable sub-questions, directs them to specialized scientist agents, and reasons about their results. The scientist agents reflect core drug R&D fields (e.g., target biology, human genetics, single cell biology, functional genomics, clinical trials, etc.). Each scientist agent has distinct system prompts, skills, and domain-specific tools available to them. This structured collaboration encourages deep data-driven analyses of primary evidence and allows for weighing of disparate sources of evidence using a unified reasoning layer to arrive at nuanced conclusions.

Agentic systems have already shown promise in scientific areas such as chemistry [14, 15] and machine learning [16, 17]. In biology, agents have been applied to automate gene editing [18], omics analysis [19, 20], nanobody design [21], and biological data analysis [22]. While these prior agentic systems focus on relatively narrow tasks, such as executing a predefined analysis workflow or designing a particular molecule, the Virtual Biotech is explicitly designed for translational reasoning across the therapeutic R&D pipeline. Compared to previous work, Virtual Biotech’s multi-agent orchestration enables the systematic integration of diverse forms of biological evidence, including clinical trial data, that have not been previously used in this context. By co-ordinating analysis over the large scale and complexity of drug discovery data, the system can cover the full breadth of the preclinical discovery workflow and answer scientific questions with more holistic, data-driven reasoning. In contrast to traditional manual workflows, the Virtual Biotech enables faster, more cross-referenced, objective, and reproducible analyses for early-stage decision-making.

In this paper, we first describe the Virtual Biotech orchestration and tooling, consisting of four scientific divisions, eleven total agents, and over 100 tools. We then demonstrate the Virtual Biotech’s data-processing capabilities by using it to analyze 55,984 clinical trials where it discovered previously unreported associations between single-cell transcriptomic features of drug targets and trial success. Next, we applied the Virtual Biotech to two real-world drug discovery scenarios (B7-H3 antibody-drug conjugate development and first-in-class mAb development against OSMR*β* in ulcerative colitis), showing that it yields useful insights into mechanisms and development strategies that complement the findings of major biopharmaceutical companies.

## 2 Results

### Overview of the Virtual Biotech

The Virtual Biotech emulates a cross-functional therapeutic research organization. It consists of four divisions of scientist agents that address key steps of the drug discovery pipeline (Figure 1A). Domain-specialized scientist agents in each division have complementary expertise and domain-specific tools that allow them to intelligently query, analyze, and integrate biomedical information across the translational spectrum for evidence-based assessments of drug discovery questions. The organization of the Virtual Biotech mirrors that of human biotech companies, which has the added benefit of enabling each agent to be audited by the corresponding human expert more easily. We created domain-specific model context protocol (MCP) servers to provide a unified programmatic interface to major biomedical databases and knowledge sources, abstracting the complexity of heterogeneous data formats and query APIs into standardized tools that agents use autonomously (Figure 1B) [23–25]. Each scientist agent is granted access to a curated subset of MCP tools aligned with its domain expertise, enabling it to gather evidence from relevant data sources while maintaining clear separation of responsibilities across the multi-agent system. This transforms static databases into interactive knowledge layers that agents dynamically analyze in response to user questions.

**Figure 1:**
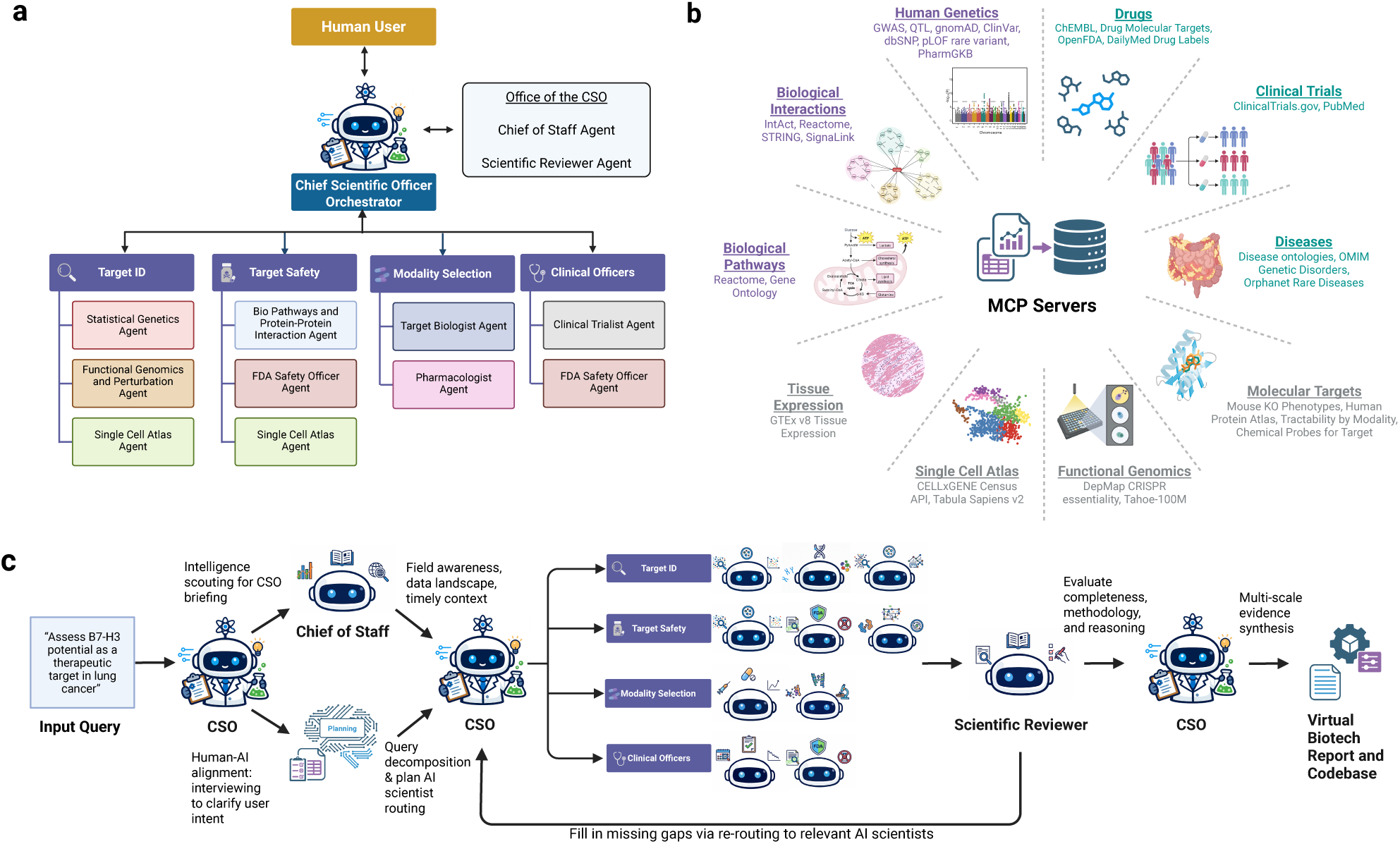
Overview of the Virtual Biotech AI agents, tools, and orchestration. (A) The multi-agent AI system organization and scientific divisions. (B) Overview of model context protocol (MCP) servers, grouped by domain area, which provide customized tools to query and analyze diverse biological and clinical data sources. (C) The workflow of the Virtual Biotech. In response to a user query, the virtual CSO first asks any relevant follow-up questions to clarify user intent before doing expensive analyses. In parallel, the chief of staff agent prepares a briefing on the topic with recent developments, data availability, and general field context. The virtual CSO then coordinates divisions of specialized scientist agents to perform analyses to answer the user’s query. Next, the scientific reviewer agent evaluates the methodology and claims from scientist agents, and upon reviewing the feedback, the CSO either coordinates revised analyses with scientist agents or synthesizes the evidence into a report.

We developed more than 100 customized MCPs and analytical tools enabling Virtual Biotech scientist agents to interrogate large-scale, heterogeneous biomedical data spanning 78,726 targets, 39,530 diseases, 14.5 million protein–protein interactions, over 3 million genetic credible sets, and more than 100 million single-cell profiles [26, 27]. The platform further integrates chemoinformatics and pharmacology resources covering 18,119 drugs, 6,332 distinct mechanisms of action, 114,270 FDA adverse event reports, and 32,783 pharmacogenomic variants, together with more than 4 billion drug perturbation gene expression measurements from Tahoe-100M [28]. In addition, ClinicalTrials.gov records are accessible through our MCP [29]. Scientist agents can further query biomedical literature through PubMed and retrieve additional targeted datasets via webbased tools.

### Scientist Divisions

The Virtual Biotech is composed of the following scientific divisions:

- **The Target Identification and Prioritization division** integrates genetic causality, functional dependency, and cellular context to validate targets.
- **The Target Safety division** integrates pathway reasoning, interaction-based collateral effects, cell type-specific expression, and historical FDA regulatory safety signals to assess potential on and off-target safety concerns.
- **The Modality Selection division** determines the optimal therapeutic modality for validated targets using structural biology, druggability assessment, and clinical precedence data.
- **The Clinical Officers division** bridges preclinical assessment with clinical reality by analyzing regulatory safety data and extracting insights from prior clinical development efforts.

### Virtual Biotech Orchestration

The Virtual Biotech is coordinated by the virtual CSO agent, which interprets user queries, clarifies research objectives, and delegates analytical tasks to scientist agents (Figure 1C). The virtual CSO oversees the overall landscape of scientific directions while not conducting any data analyses itself. Through its system prompt, it was designed to know (1) what questions to ask, (2) who to ask those questions to, and (3) how to synthesize multi-scale evidence. Upon receiving a user’s query and before initiating scientist work, the CSO initiates its own strategic orientation. The virtual CSO calls upon its chief of staff agent to generate a briefing on general field awareness, data landscape, and recent developments, which the chief of staff agent obtains by reviewing Virtual Biotech infrastructure and performing web searches. This briefing helps the CSO frame the analysis, set proper feasibility expectations for the user, and identify key questions to prioritize. In parallel, the CSO interviews the user to clarify the intent of the user and increase the alignment of human-AI collaboration [30, 31].

With the chief of staff briefing and the clarified intent of the user, the virtual CSO is then properly oriented for the task. It then performs task decomposition to break the query into sub-tasks that are intelligently routed to the relevant scientist agents. The CSO can engage all relevant divisions either simultaneously or one after another in defined sequences (Figure 1C). This orchestration pattern is task-dependent and the CSO has the authority to control when and how information flows between divisions.

After the scientist agents complete their analyses, the scientific reviewer agent assesses their outputs based on three criteria: how well they address the user’s original question, the strength of the evidence supporting their conclusions, and the overall thoroughness of the analysis (Figure 1C, Supplementary Note II). If gaps or unsubstantiated claims are identified by the scientific reviewer agent, the CSO re-delegates to relevant scientist agents with the reviewer agent’s feedback for iterative refinement. This orchestration pattern separates strategic reasoning from domain-specific analyses, enabling the CSO to synthesize findings across divisions into cohesive recommendations while each scientist agent focuses on its domain of expertise.

We developed a user interface for the Virtual Biotech that displays the agents’ reasoning process, indicates which tools are currently in use, and allows users to download the corresponding data, reproducible code, and reports generated during each session (Figure S1).

### Virtual Biotech connects preclinical biology with clinical outcomes to identify novel drug target features associated with trial success

Drug discovery programs face substantial attrition, with lack of efficacy and safety concerns accounting for approximately 79% of clinical trial failures [32]. Prior work has established that targets supported by human genetic evidence are significantly more likely to progress through clinical development, with genetically supported drugs over twice as likely to reach approval and genetic evidence underlying two-thirds of recent FDA-approved therapeutics [33–35]. To identify opportunities for expanding on these results, we provided Virtual Biotech with this earlier work and instructed the system to critically evaluate its own infrastructure and recommend new directions to expand this analysis [34]. Following discussions with its scientific divisions, the virtual CSO suggested leveraging its clinical trialist agent to systematically enrich the existing Open Targets clinical trial dataset—which currently encompasses 55,984 trials—with additional outcome annotations. It also proposed using its single-cell and functional genomics agents to design genomic features to be tested for association with trial outcomes following the approach of Razuvayevskaya et al. [27, 34, 36]. Generating outcome annotations with the clinical trialist agent was necessary because the original Open Targets dataset did not include endpoint data, and the corresponding ClinicalTrials.gov records frequently have incomplete, inconsistently reported, or free-text-only outcomes, limiting their utility for downstream analyses [37, 38]. Curation by clinical trialist agents was therefore required to standardize these data and recover missing or discrepant outcomes from additional sources such as publications or press releases.

#### Massively parallel, source-tracked extraction of trial outcomes

The clinical trialist agent proposed key outcome variables for large-scale extraction, including trial phase progression, primary and secondary endpoint results, and adverse event rates. It then designed statistical analyses to test whether genetic evidence and single-cell genomic characteristics of drug targets were associated with these outcomes. However, the aforementioned outcome variables are not available in the Open Targets dataset and were frequently missing or inconsistently reported in ClinicalTrials.gov. To address this, the clinical trialist agent developed a systematic protocol to obtain trial results from sources (e.g. publications, press releases) and to produce standardized, harmonized outcome annotations.

To enable high-throughput curation at scale, the Virtual Biotech created and dispatched 37,075 clinical trialist agents in parallel to extract outcome data from multiple sources for Phase II and III trials in the Open Targets dataset (Figure 2A). At this scale, comprehensive curation is impractical without parallelization, necessitating a multi-agent architecture that can execute data curation in a distributed manner. Each clinical trialist agent followed a three-tier evidence cascade for its assigned National Clinical Trial (NCT) ID: it first systematically queried the ClinicalTrials.gov database, then searched PubMed for publications containing trial results when primary data were incomplete, and finally consulted online press releases and regulatory announcements for trials still missing required data fields (Supplementary Note I) [29]. For reproducibility and transparency, agents recorded the source of information for each annotation. For Phase I trials, instead of manual curation by an agent, success was algorithmically defined as progression to Phase II, since Phase I trials are typically designed to assess safety and dosing rather than efficacy. Phase I trials were classified as having progressed to Phase II if there existed at least one Phase 2 trial which tested the same set of drugs for the same indications, with a subsequent or concurrent trial start date.

**Figure 2:**
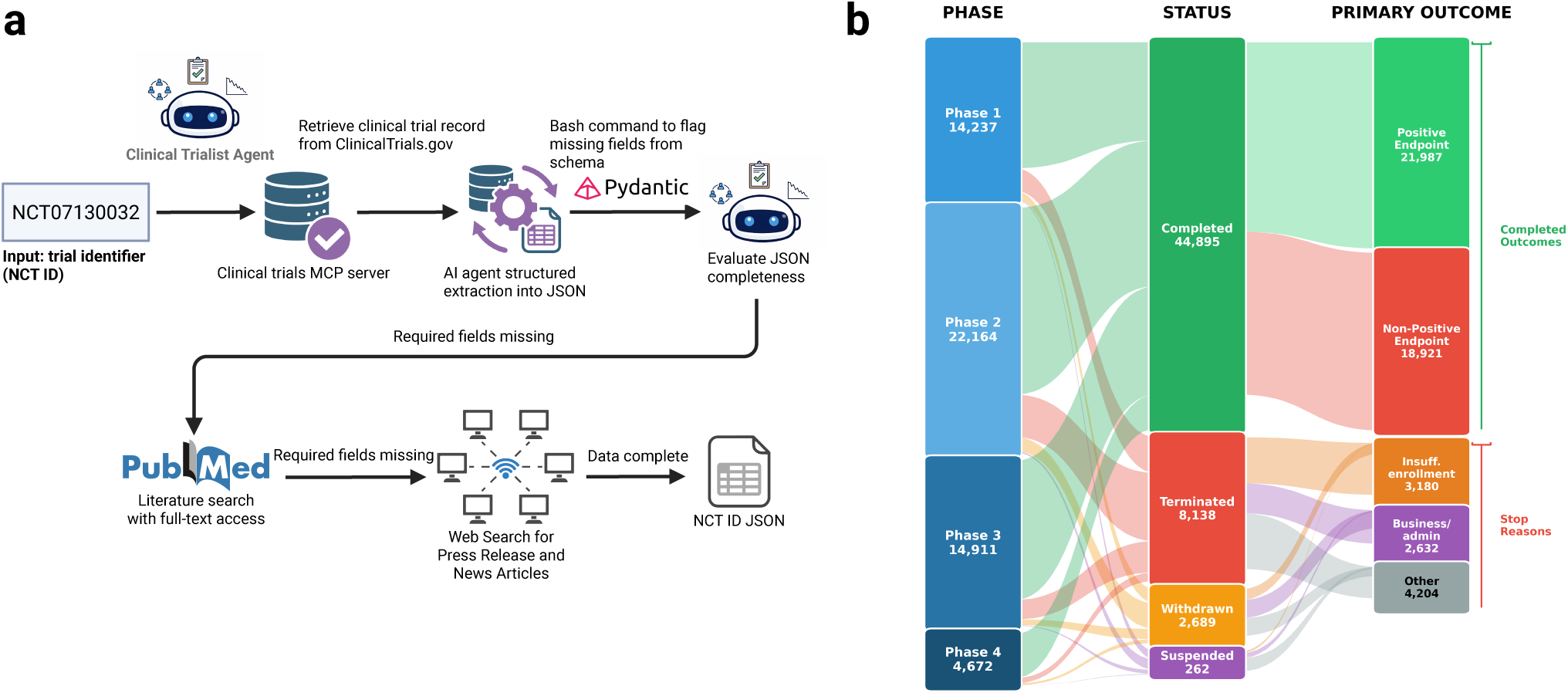
Large-scale clinical trial data extraction using the Virtual Biotech. (A) Overview of the data extraction and curation process. For each National Clinical Trial (NCT) ID, the clinical trialist agent uses tool calls to ClinicalTrials.gov, PubMed, and the web to gather information about the trial. After reasoning about its findings, the agent enters information into a structured JSON ontology for downstream analysis. (B) Sankey plot of the curated clinical trial dataset (n=55,984) showing trial phase, trial status, and outcomes.

We manually evaluated the quality of agent annotations by reviewing 100 randomly selected clinical trials and measuring the concordance between the human reviewer and the agent. For both primary and secondary endpoints, trials were classified into two groups based on whether or not there was sufficient evidence supporting a positive trial endpoint. Not all outcomes were applicable to each trial, e.g., primary endpoints were not counted for trials stopped early. Agreement between human reviewer and the agent was 89.7% (78/87) for primary endpoints, 83.9% (73/87) for secondary endpoints, and 92.4% (61/66 that reported exact statistics) for adverse event (AE) rates.

We further found a 85.3% agreement in primary endpoints across 7,278 overlapping trials between the Virtual Biotech dataset and the manually curated Therapeutic Data Commons (TDC) Trial Outcome Prediction dataset [39]. Among the trials where the agent’s annotations differed from manual review or TDC data, the differences were largely confined to intrinsically ambiguous cases: studies with multiple experimental arms in which some arms achieved the primary endpoint while others did not; trials where the primary endpoint failed to reach conventional statistical significance (e.g., p=0.051) despite supportive secondary or longer-term outcomes; and single-arm or dosage-based study designs that do not map cleanly onto a simple success/failure dichotomous outcome. These edge cases indicate intrinsic ambiguity in reducing complex, multi-endpoint results to discrete classes, rather than systematic mistakes by either annotation source.

Taken together, these agreement metrics demonstrate that the agent assigned labels to trials reliably and that the resulting annotations are suitable for subsequent analyses. We release the resulting dataset, forming one of the largest and most comprehensive publicly available datasets to date of clinical trial outcomes linked to drug targets. Compared to existing resources, the Virtual Biotech dataset is unique in its scale and source-tracked annotations that harmonize endpoint outcomes and adverse event rates across reporting sources.

#### Translational feature engineering with cell atlases and perturbation signatures

To derive single-cell expression features of drug targets, the single cell agent proposed and implemented a comprehensive feature extraction framework for Tabula Sapiens v2, a healthy human reference atlas of 27 tissues [40]. The agent designed features organized into three categories: cell-type specificity metrics, expression heterogeneity measures, and drug perturbation response signatures. For specificity, the agent proposed the tau cell-type specificity index, where values approaching zero indicate ubiquitous expression while values approaching one indicate expression restricted to a single cell type [41]. To capture expression heterogeneity within cell populations, the agent proposed a bimodality coefficient calculated from the skewness and kurtosis of expression distributions among expressing cells, where higher values may indicate "on/off" expression patterns within a tissue [42, 43]. The bimodality coefficient is widely used in psychology and social sciences but less common in single-cell analysis, reflecting cross-domain knowledge transfer [44–46]. We found that these features are moderately correlated (Pearson *ρ* = 0.54), suggesting that they provide complementary information.

Separately, the functional genomics agent proposed to leverage Tahoe-100M [28], a large-scale perturbational transcriptomic resource, to construct six hallmark expression signatures: apoptosis induction, proliferation suppression, cell cycle arrest, DNA damage response, stress response, and resistance mechanisms [28]. The agent reasoned that these six signatures could demonstrate broadly applicable mechanisms of drug sensitivity and resistance reflected in perturbational expression profiles. It then designed efficacy scores for each hallmark using log-fold changes of genes from differential expression analysis comparing drug-perturbed to unperturbed cells to quantify the degree to which drug treatment induces expression changes in canonical genes for each signature (see Extended Methods for details).

#### Virtual Biotech-driven analysis reveals genomic features associated with trial success

The final integrated dataset comprised 55,984 clinical trials spanning Phase I (n=14,237), Phase II (n=22,164), Phase III (n=14,911), and Phase IV (n=4,672) (Figure 2B). Trial-level outcome annotations were linked to target-level molecular features extracted by the Virtual Biotech and to genetic evidence scores from Open Targets.

The clinical trialist agent first successfully replicated the genetic evidence results reported by Razuvayevskaya et al., finding that genetic support is associated with trial progression and also protective of early trial stoppage (Figure S2A) [34]. Using the additional trial outcomes curated by the Virtual Biotech, it also found that trials which achieved primary and secondary endpoints were enriched with genetic evidence (Figure S2A). Following the approach from the analogous study on genetic evidence, the clinical trialist agent then evaluated the association between single-cell transcriptomic features of drug targets and trial outcomes [33, 34].

Both tau cell-type specificity and expression bimodality score had significant associations with trial progression through clinical development phases (Figure 3A). Following binarization of targets based on their tau metric into cell-type-specific versus broadly expressed classes (see Extended Methods for details), the clinical trialist agent found that drugs acting on cell-type-specific targets were 48% more likely to ever reach Phase IV. Lower bimodality scores were also significantly associated with trials which were terminated (OR = 0.81; 95% CI: 0.79–0.83), withdrawn (OR = 0.88; 95% CI: 0.85–0.92), or suspended (OR = 0.70; 95% CI: 0.62–0.80) (Figure 3A).

**Figure 3:**
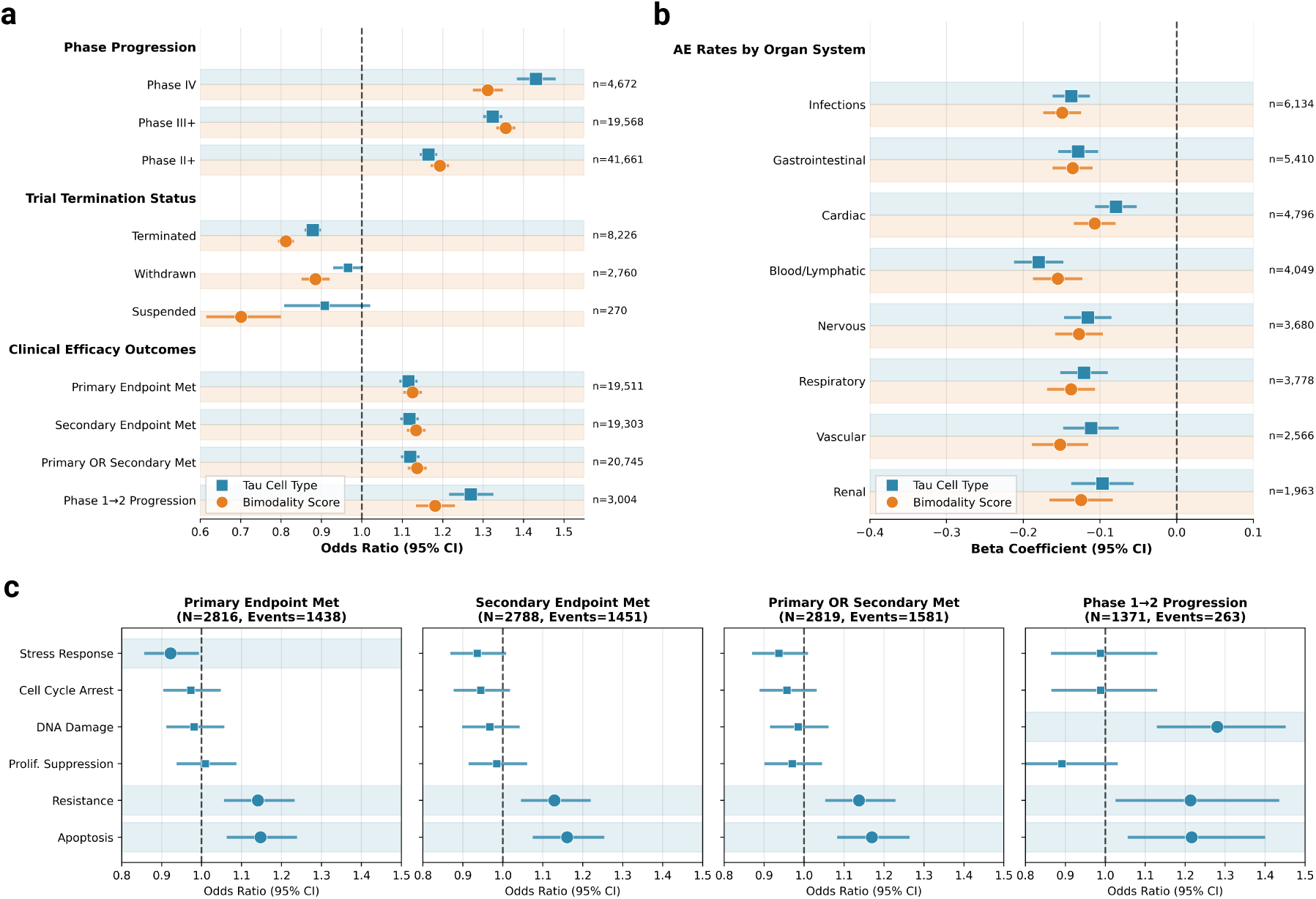
Preclinical single-cell and perturbation atlas features of drug targets associate with key clinical trial outcomes. (A) Associations between single cell-derived features of the drug target and phase progression, trial stoppage, and efficacy outcomes. Plot shows the standardized odds ratio (all features normalized to have a standard deviation of 1) with 95% confidence intervals. (B) Associations between single cell-derived features and serious adverse event rates grouped by organ system. Forest plot shows beta regression coefficients with 95% confidence intervals. (C) Associations of in vitro transcriptomic signature responses to drug perturbations and efficacy outcomes in oncology trials. Forest plot shows odds ratio with 95% confidence intervals.

Targets with higher cell type specificity were more likely to meet defined primary endpoints (OR = 1.11; 95% CI: 1.09–1.14), secondary endpoints (OR = 1.12; 95% CI: 1.10–1.14), and successful Phase I to Phase II progression (OR = 1.27; 95% CI: 1.22–1.33) (Figure 3A). Drugs with cell-typespecific targets were 40% more likely to progress from Phase I to Phase II. Similarly, targets with a higher expression bimodality score were more likely to achieve a positive primary endpoint (OR = 1.13; 95% CI: 1.10–1.15) a positive secondary endpoint (OR = 1.13; 95% CI: 1.11–1.16), and successful Phase I to Phase II progression (OR = 1.18; 95% CI: 1.13–1.23).

The Virtual Biotech further found that trials with targets that have a higher cell type specificity and expression bimodality score reported significantly lower infectious, gastrointestinal, cardiac, hematologic, nervous, respiratory, vascular, and renal adverse event rates (Figure 3B). Trials testing cell-type-specific targets had on average 32% lower adverse event rates across these organ system groups. These findings suggest that targets with more heterogeneous and cell-type-restricted expression patterns may offer improved safety profiles.

Next, we demonstrate that these findings are robust to confounders. To assess whether the observed associations could arise by chance or through confounding in the complex trial-to-target mapping, we performed permutation testing over 1,000 iterations, confirming that all findings remained significant against the empirical permuted null (Figure S3A,B). To control for potential confounding between disease indications and between drug modalities, we estimated ORs using mixed-effects models adjusted for the phase and enrollment year, with the trial’s drug modality and disease indication area as random effects. The adjusted odds ratios for cell type specificity and expression bimodality score remained significant across mixed-effects models (Figure S3C,D).

Another key question is whether single-cell transcriptomic features of targets provide independent value beyond established human genetic evidence linking targets to disease. Therefore, we replicated our findings adjusting for genetic evidence. Specifically, we found that both cell type specificity and expression bimodality score retained all significant associations with negligible decreases in effect sizes after adjustment (Figure S4A,B). We further estimated ORs for single-cell features among trials with no genetic evidence (74.6% of trials) and found that single-cell features had similar or even stronger effect sizes in this subset (Figure S4C,D). Together, these results indicate that single-cell information captures additional biological signal associated with clinical trial outcomes beyond genetic evidence alone. These findings are among the first to demonstrate that single-cell features can be leveraged to assess targets within drug discovery programs. A recent report by Dann et al., which focused only on phase progression, found that cell-type-restricted targets were associated with early progression through phases [47]. They did not study association with safety or whether the trial endpoints were met.

Beyond cell-type specificity and bimodality, the Virtual Biotech examined whether features of hallmark pathways derived from drug perturbation responses of investigational or FDA-approved molecules in the Tahoe-100M atlas were associated trial efficacy. The functional genomics agent mapped a total of 3,345 unique drug-cell line combinations to 7,920 oncology trials which tested the identical drug and cancer histology indication. Several hallmark scores showed significant associations with trials that achieved stated endpoints in phase-adjusted analyses. The apoptosis score was positively associated with primary endpoint success (OR = 1.15; 95% CI: 1.06–1.24), suggesting that drugs inducing stronger pro-apoptotic transcriptional signatures in vitro for the relevant cell lines were more likely to achieve primary endpoint success in its corresponding clinical trial (Figure 3C). The resistance score was similarly associated with increased odds of achieving primary endpoint (OR = 1.14; 95% CI: 1.06–1.23), consistent with the selection of drugs that overcome cellular defense mechanisms. Similar patterns emerged for secondary endpoint success. For Phase I to Phase II progression, the DNA damage score showed the strongest association (OR = 1.28; 95% CI: 1.13–1.45), followed by apoptosis (OR = 1.22; 95% CI: 1.06–1.40), resistance (OR = 1.21; 95% CI: 1.03–1.43). The prominence of DNA damage response in early-phase progression may reflect the importance of demonstrating clear pharmacodynamic activity in Phase 1 studies. As one of the first analyses to link drug-perturbation atlas single-cell features to trial outcomes at scale across thousands of drug–histology trials, these results highlight the utility of perturbation biology in assessing downstream clinical efficacy.

### Virtual Biotech research builds preclinical rationale for B7-H3 antibody-drug conjugate development in lung cancers

To demonstrate the Virtual Biotech’s capacity to rapidly generate rigorous and reproducible evidence to support therapeutic development, we tasked the system to assess the therapeutic potential of B7-H3 (also known as CD276) in lung adenocarcinoma (LUAD) and small cell lung cancer (SCLC). B7-H3 is a transmembrane immune checkpoint protein that has been reported to be highly expressed in cancer cells in a wide range of solid tumors, including lung cancers [48]. Its receptor is unknown, and its role in cancer remains unclear, though it likely has both immunologic and non-immunologic functions [49]. To evaluate its potential as a therapeutic target, the Virtual Biotech analyzed and integrated evidence from human genetics, target biology, single-cell transcriptomics, spatial transcriptomics, biomarker-stratified survival analyses, and prior clinical trials.

The statistical genetics agent first evaluated germline genetic support for B7-H3 as a lung cancer target. No significant associations were identified at the B7-H3 locus for lung cancer (Figure 4A). Enhancer-to-gene (E2G) analysis in lung tissue with ENCODE identified 41 high-confidence regulatory elements at the B7-H3 locus, consistent with strong transcriptional activity and open chromatin in normal lung. Constraint metrics showed a predicted loss-of-function observed-to-expected ratio of 0.67, suggesting moderate tolerance to inactivating variants. Collectively, the agent concluded that germline genetic evidence for B7-H3 as a lung cancer target is weak, though strong regulatory activity in lung tissue supports its expression and potential functional relevance. Despite this, the virtual CSO correctly interpreted the absence of GWAS signal as non-disqualifying. Upon receiving the results, it noted that for potential immune checkpoint targets, therapeutic rationale derives from somatic tumor overexpression rather than germline genetic risk variants. This reasoning distinguished B7-H3 from targets where genetic evidence is essential for validation, prompting the virtual CSO to instruct its single cell agent to perform analyses for more clarity.

**Figure 4:**
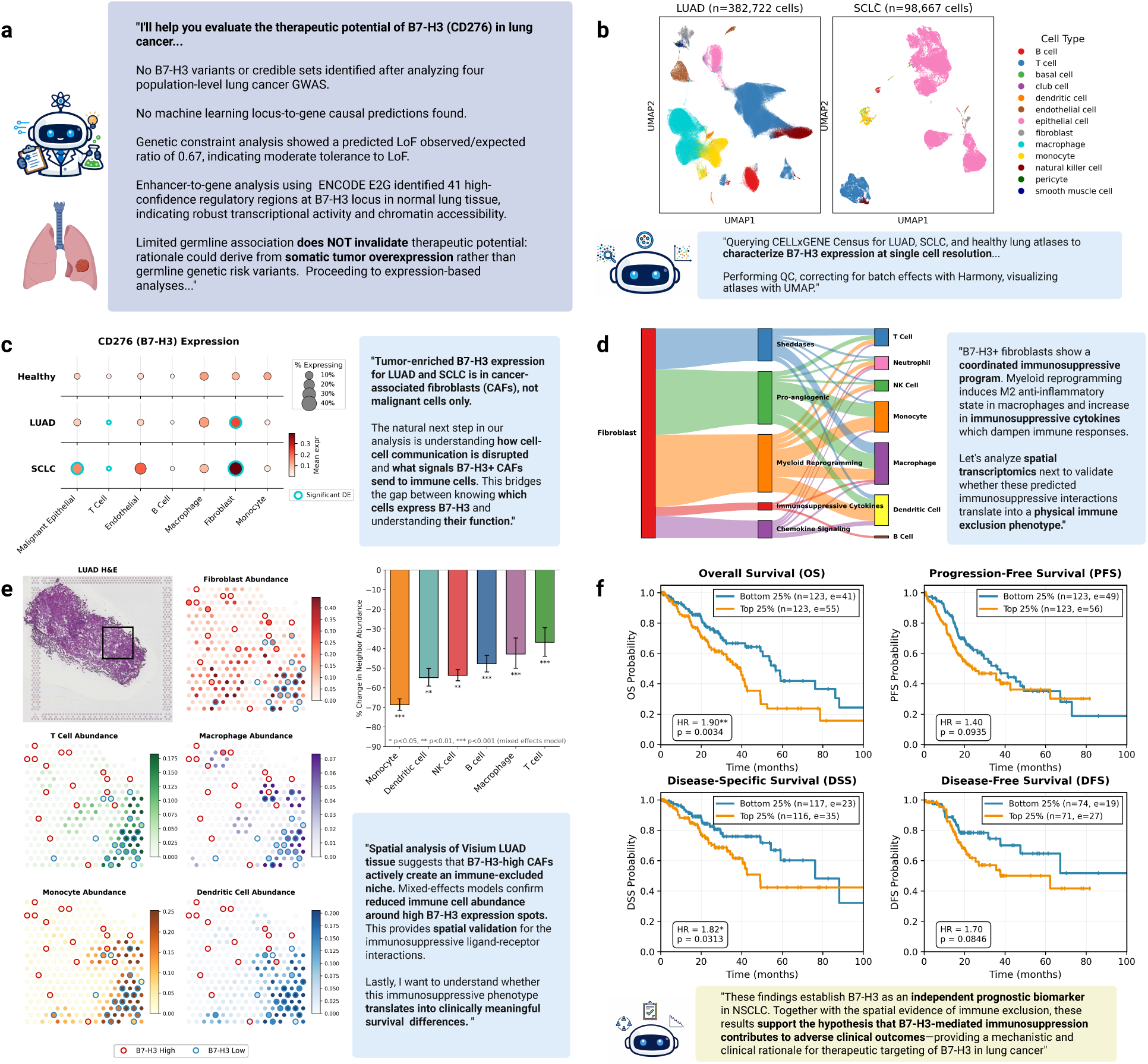
Virtual Biotech evaluates B7-H3 (CD276) as therapeutic target in lung cancer. (A) Summary from statistical genetics agent of germline genetic findings for B7-H3 in lung cancer (B) UMAP of lung adenocarcinoma (LUAD) and small cell lung cancer (SCLC) single cell atlases prepared by the single cell atlas agent. (C) Differential expression (DE) analysis of B7-H3 among cell types by the single cell atlas agent identifies significantly increased expression in fibroblasts. (D) Ensemble cell-cell communication inference by the single cell agent reveals unique signaling pathways found in B7-H3+ cancer-associated fibroblasts in SCLC (shown here) and LUAD (see Supplemental Figure S5). (E) Spatial transcriptomic analyses of LUAD primary-site tissue identifies immune-excluded niches in B7-H3 expressing spots relative to B7-H3 non-expressing spots. Bar plot quantifies the percent change in neighbor immune abundance between top and bottom quartile of B7-H3-expressing spots across 12 LUAD tumor samples. Significance was assessed using a mixed-effects model adjusting for total UMI count, fibroblast abundance, and malignant epithelial abundance fixed effects, with random effects at the sample-level. (F) Survival analysis of the NSCLC cohort from The Cancer Genome Atlas project identifies B7-H3 as a prognostic biomarker of disease-specific survival.

To characterize B7-H3 expression at cellular resolution, the single cell agent queried and downloaded single-cell atlases from CELLxGENE Census, comprising 92,061 cells from 9 donors in the SCLC atlas, 337,002 cells from 69 donors in the LUAD atlas, and 86,478 cells from lung tissue of 26 healthy donors (Figure 4B) [50]. The agent then conducted cell-type-resolved differential expression analysis with pseudobulk aggregation, revealing that B7-H3 upregulation was not uniform across cell types but rather concentrated in fibroblasts in both SCLC (log2FC = 1.94, FDR P-value = 3.0 × 10^−6^) and LUAD (log2FC = 1.46, FDR P-value = 1.1 × 10^−7^) (Figure 4C). T cells showed a significant downregulation of B7-H3 in both types of cancer (SCLC log2FC: -1.822, FDR P-value = 5.5 × 10^−3^; LUAD log2FC: -1.924, FDR P-value = 6.0 × 10^−8^). Based on these findings, the single cell agent hypothesized that B7-H3-high fibroblast cells might actively shape the immunosuppressive microenvironment. To investigate this, it performed a ligand–receptor interaction analysis with LIANA+ to identify differences in signaling between fibroblasts whose B7-H3 expression fell within the highest vs. lowest quartiles [51]. This analysis identified 152 significant interactions in SCLC and 56 interactions in LUAD that were specific to B7-H3-high fibroblasts communicating with immune cells (Figure 4D). Functional characterization of these interactions by the single cell agent identified five predominant signaling classes: sheddases that cleave activating ligands from immune cell surfaces, myeloid-reprogramming mediators that drive macrophage polarization toward immunosuppressive phenotypes, pro-angiogenic factors that remodel the tumor vasculature, immunosuppressive cytokines that engage inhibitory receptors, and chemokine pathways that retain immune cells within the fibroblast niche. Together, these findings raised the possibility that B7-H3-high stromal cells leverage immunomodulatory mechanisms in addition to the canonical checkpoint activity of B7-H3 itself.

At this point, the virtual CSO dispatched its scientific reviewer agent. The reviewer agent noted that the preceding ligand-receptor analysis revealed that B7-H3-high fibroblasts engage in distinct immunosuppressive communication patterns compared to B7-H3-low cells, but this was inferred from dissociated single-cell data that inherently lose spatial tissue context. With this, the virtual CSO then reasoned that validating whether these predicted immunosuppressive interactions translate into a physical immune exclusion phenotype required spatial profiling. It then dispatched the single cell agent again to analyze a Visium dataset comprising 12 lung tissue samples from patients diagnosed with LUAD, totaling approximately 65,000 spatially-resolved spots after quality control [52]. To infer cell type composition at each spot, the agent applied Cell2Location using the prior LUAD single-cell atlas, imputing proportions for 13 cell types [53]. The agent subsequently categorized Visium spots on each tissue slide into top or bottom quartiles based on their B7-H3 expression levels. It then implemented a mixed-effects regression model to assess whether B7-H3-high spots showed altered immune cell compositions within their immediate spatial neighborhood (k = 6 nearest spots) adjusting for sequencing quality, fibroblast content, malignant epithelial cell abundance, and sample-to-sample variability through random effects (Figure 4E). The single cell agent’s mixed-effects model found that B7-H3-high spots exhibited significant depletion of neighboring immune cells such as T cells, macrophages, monocytes, and dendritic cells (Figure 4E). This confirmed that B7-H3 expression spatially correlates with an immune-excluded tumor microenvironment, further supporting the hypothesis of an immunosuppressive phenotype predicted from single-cell ligand-receptor analysis and providing functional evidence supporting B7-H3 as an checkpoint target in lung cancer.

Next, the virtual CSO wanted to understand whether this immunosuppressive profile corresponded to clinically significant differences in patient survival. To address this, the virtual CSO tasked the clinical trialist agent with performing a biomarker-stratified survival analysis on the TCGA PanCancer Atlas LUAD cohort, utilizing the cBioPortal MCP tools for data access [54, 55]. This dataset includes 566 patients with RNA-seq gene expression profiles and detailed survival outcomes. To maximize separation between biologically distinct groups, the clinical trialist agent stratified patients by B7-H3 expression quartiles, comparing patients in the top and bottom quartiles based on their B7-H3 expression. In a Cox proportional hazards models adjusted for age, cancer stage, and sex, elevated B7-H3 expression was significantly linked to worse overall survival (HR = 1.62, 95% CI: 1.05–2.48, *p* = 0.028) and worse disease-specific survival (HR = 1.82, 95% CI: 1.06–3.15, *p* = 0.031) (Figure 4F). Based on these results, the virtual CSO proposed that B7-H3 acts as a prognostic biomarker for DSS in LUAD and, when interpreted together with spatial evidence of immune exclusion, bolsters the notion that B7-H3–mediated immunosuppression drives unfavorable clinical outcomes—thus providing both mechanistic and clinical rationale for therapeutically targeting B7-H3 in lung cancer.

Finally, to determine the best therapeutic modality for targeting B7-H3, the virtual CSO dispatched the target biologist agent. This agent used MCP tools to retrieve Human Protein Atlas summaries of B7-H3 that confirmed cell surface expression with subcellular vesicle localization [56]. Together with single-cell findings showing that B7-H3 is predominantly overexpressed within tumor microenvironment (TME) fibroblasts and not only on malignant cells, the virtual CSO concluded that an antibody–drug conjugate (ADC) targeting B7-H3 would be a promising therapeutic approach. The virtual CSO supported its position by noting that an ADC’s effectiveness can operate through several mechanisms consistent with its findings: targeted delivery of a cytotoxic payload to cancer cells that express B7-H3, alteration of the TME through inhibition of B7-H3–mediated immunosuppressive checkpoint interactions, and ADC-driven bystander killing of adjacent cells that lack B7-H3 expression. Small molecule and proteolysis targeting chimera (PROTAC) approaches were de-prioritized after the pharmacologist agent’s tractability analysis identified no druggable protein pockets and no known chemical binders for B7-H3. The clinical trialist agent was subsequently instructed by the CSO to query ChEMBL to assess the competitive landscape and uncover clinical precedents. Through this analysis, it determined that the ADC modality has been validated in metastatic HER2-positive breast cancer with trastuzumab deruxtecan, which demonstrated significant efficacy and obtained FDA approval. The search also showed that the naked anti–B7-H3 antibody enoblituzumab is in ongoing Phase 2 development for prostate cancer, and that no B7-H3–directed therapies have yet been approved, highlighting both a clear opportunity for differentiation and a remaining unmet medical need.

The Virtual Biotech conducted this case study without access to web search or fetch tools to prevent information leakage, and its underlying language model had a training knowledge cutoff of January 2025. Importantly, existing literature emphasized B7-H3’s expression on cancer cells and canonical checkpoint-associated functions, motivating antibody- and ADC-based programs largely through a cancer-cell–centric lens [48, 49]. In contrast, the Virtual Biotech independently integrated orthogonal evidence modalities to nominate a less-explored but mechanistically coherent hypothesis in lung cancers: that B7-H3 is more strongly overexpressed in cancer-associated fibroblasts, spatially associates with immune exclusion, and stratifies poor clinical outcomes, an interpretation that is not directly identified from any single dataset or previous publication. Furthermore, in August 2025, the FDA granted Breakthrough Therapy Designation to ifinatamab deruxtecan, a first-in-class B7-H3–targeted ADC that demonstrated a 48.2% objective response rate in a phase II trial in SCLC [57–59]. As the Virtual Biotech did not have access to this information— it came after the model’s knowledge cutoff—the trial’s results provides independent external support for the system’s mechanism-driven therapeutic design.

This systematic and data-driven analysis covering genetics, single-cell and spatial transcriptomics, clinicogenomics, and modality assessment was completed for a total cost of $46.00 in Anthropic API credits. It showed how the Virtual Biotech can accelerate target identification and validation, a typically manual process performed by cross-functional drug discovery teams.

### Virtual Biotech revises development strategy for terminated Phase II trial testing first-in-class mAb against OSMR*β* in ulcerative colitis

We subsequently used the Virtual Biotech to analyze instances of late-stage clinical development failure. We focused on the Phase II MOONGLOW study (NCT06137183), a trial in ulcerative colitis (UC) evaluating the oncostatin M receptor subunit OSMR*β* as a therapeutic target, which was terminated in August 2025 [60]. OSMR*β* is a cytokine receptor subunit that heterodimerizes with gp130 or the IL-31 receptor-*α* to mediate oncostatin M and IL-31 signaling—pathways implicated in inflammation and fibrosis, ultimately activating canonical JAK-STAT cascades that include STAT3 [61, 62]. Vixarelimab is a first-in-class, human monoclonal antibody engineered to block OSMR*β*-mediated signaling while preserving the LIFR pathway [60]. In NCT06137183, adults with moderate-to-severe UC were randomized to receive vixarelimab or placebo with an active-treatment extension. However, the study was stopped after 79 participants were enrolled because an interim futility analysis indicated the primary endpoint was unlikely to be achieved [60]. No serious adverse events were reported among the participants [60].

We instructed the Virtual Biotech to review the target biology and trial characteristics to generate insights for why the trial failed. First, the statistical genetics agent evaluated the genetic support for OSMR*β* as a UC target. The agent identified rs395157 in the OSMR locus on chromosome 5 (OR = 1.06, p = 2.68 × 10^−10^) from a multi-ancestry meta-analysis with 23,252 cases and 352,256 controls [63]. Fine-mapping using SuSiE-inf assigned a posterior probability of 99.97% to rs395157, indicating confidence of the causal signal [63]. Locus-to-gene (L2G) prediction—which integrates functional genomic annotations, expression quantitative trait loci, and chromatin accessibility—assigned OSMR an L2G score of 0.970, indicating high confidence that OSMR is the causal gene at this locus. Next, the single cell atlas agent evaluated OSMR dysregulation in UC tissue by retrieving a UC disease atlas from CELLxGENE which included 9 UC donors and 46 healthy donors [50, 64]. Pseudobulk differential expression analysis in 25 cell types revealed no significant differences in OSMR expression between UC and healthy tissue.

In light of the mixed pre-clinical findings, the virtual CSO instructed the clinical trialist agent to analyze the NCT06137183 trial design to identify possible reasons of failure (Figure 5A). The clinical trialist agent assessed key trial features from the ClinicalTrials.gov entry, which included the full study protocol (Protocol GA44839, Version 2 PDF) [60]. In its analysis, the agent focused on the inclusion and exclusion criteria. First, it observed that the protocol excluded UC patients who showed an inadequate response to previous JAK inhibitor treatment. The clinical trialist agent noted this aligned with the bio-pathway agent’s report that OSMR acts upstream via the JAK–STAT pathway. Therefore, including patients who had already not responded to JAK inhibitors would likely select for patients in whom this common downstream signaling route is ineffective. Separately, the clinical trialist agent observed that the inclusion criteria required a treatment-refractory study population, which specified the enrollment of patients with moderate-to-severe UC who had shown poor response to conventional therapies (corticosteroids and/or immunosuppressants) or advanced therapies (TNF inhibitors, anti-integrin, or anti–interleukin). The agent therefore flagged the lack of OSMR-based stratification as a notable design limitation: without prospectively selecting an OSMR-high disease endotype, the trial risks testing an on-target mechanism in a population where many participants may not have the target pathway active at baseline. This gap directly motivated the agent to investigate the biomarker potential of OSMR in

**Figure 5:**
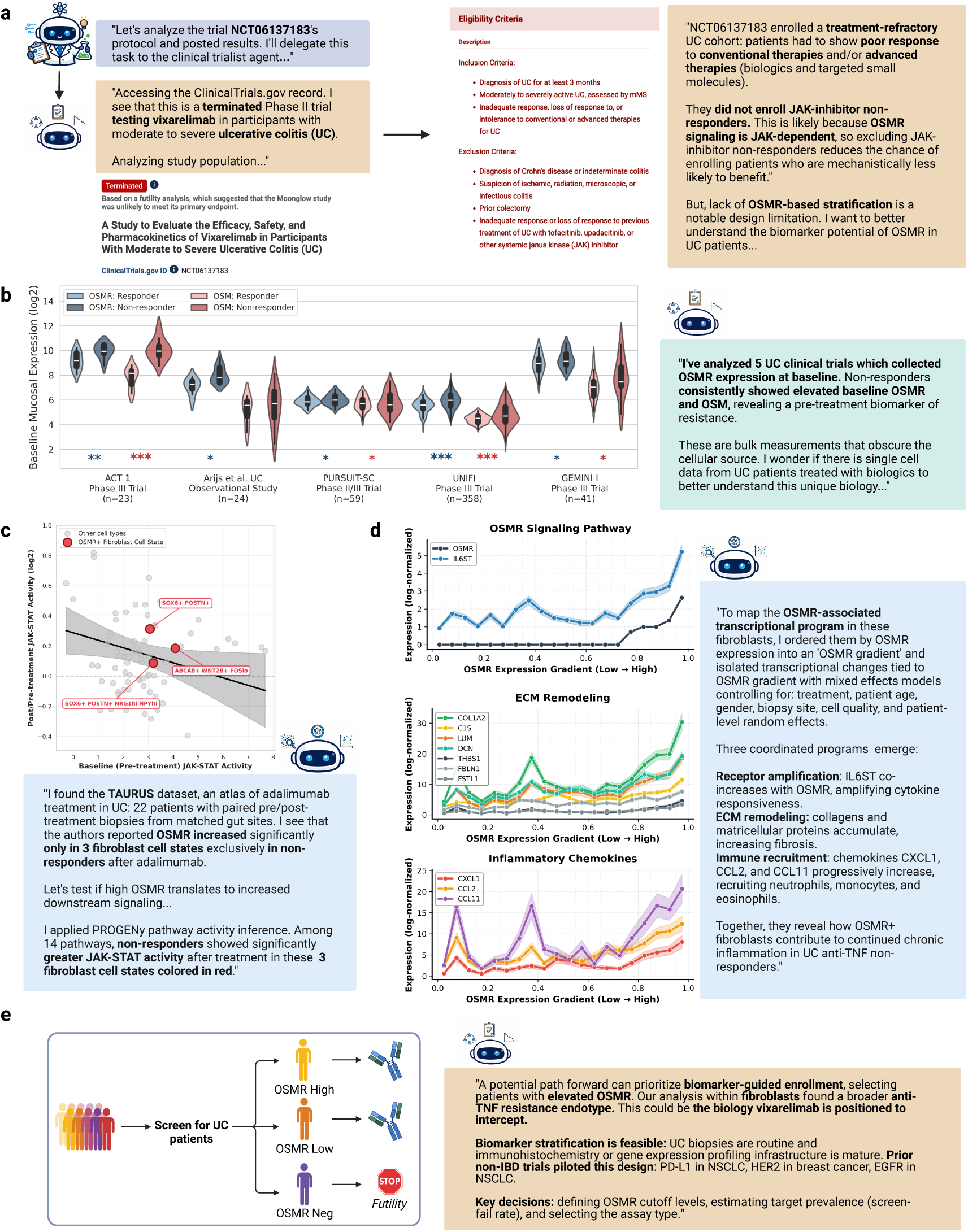
Retrospective evaluation of terminated Phase II trial protocol by the Virtual Biotech leads to refined development strategy for an OSMR*β*-directed therapy in ulcerative colitis (UC). (A) Agent summary of the Phase II UC trial (NCT06137183) investigating vixarelimab. The clinical trialist agent examined the trial protocol and reasoned through the study population. (B) Cross-trial comparison of baseline OSMR and OSM expression showing consistently higher pre-treatment levels in non-responders versus responders across multiple UC biologic trials. Significance levels from Wilcoxon rank-sum tests: *p < 0.05, **p < 0.01, ***p < 0.001. (C) Analysis of single-cell ulcerative colitis atlas of anti-TNF treatment using PROGENy-inferred JAK-STAT pathway activity. (D) OSMR expression gradient analysis within fibroblasts extract OSMR-linked transcriptional programs, revealing coordinated increases consistent with receptor amplification, ECM remodeling, and immune recruitment chemokines along the gradient. (E) Agent-proposed patients with UC.

First, the clinical trialist agent systematically searched and collected baseline gene expression data from five independent clinical trials of UC patients treated with biologics [65–69]. Notably, the agent identified trials testing biologics with diverse mechanisms to enable comparisons. These trials tested unique therapeutic mechanisms: (1) anti-TNF, (2) anti-IL12/23 blockade, and (3) anti-integrin therapy [65–69]. These trials collected colonic mucosal biopsies at baseline and measured gene expression using microarrays. Treatment response was determined by each trial’s own protocol using clinical and endoscopic endpoints. The clinical trialist-designed analyses found that across all five trials spanning three therapeutic mechanisms, biologic non-responders consistently showed significantly higher baseline OSMR expression than responders (Wilcoxon rank-sum test; anti-TNF: ACT 1 *p* = 0.006, Arijs et al. observational study *p* = 0.019, PURSUIT-SC *p* = 0.047; anti-IL12/23: UNIFI *p <* 0.0001; anti-integrin: GEMINI I *p* = 0.049). Similar patterns were observed for OSM expression (Figure 5B).

Motivated by this distinctive baseline OSMR expression signature and aiming to dissect it at both cell-type and temporal resolution, the single cell agent further identified the TAURUS dataset, a longitudinal single-cell transcriptomic atlas from paired gut biopsies collected before and after adalimumab (anti-TNF) treatment [70]. The agent focused on 22 UC patients (485,868 cells), of whom 8 achieved remission. From the supplementary materials of the original longitudinal expression analysis, it identified that OSMR expression increased significantly only in three fibroblast cell states exclusively in non-responders after treatment. The single cell agent next applied PROGENy pathway activity inference with TAURUS data to assess signaling dynamics in these fibroblasts of non-responders across 14 pathways (Figure 5C). These findings showed that JAK-STAT pathway activity was markedly elevated after anti-TNF treatment across these fibroblast cell states, shedding light on the functional implications of increased OSMR expression in non-responders. In these same fibroblasts, the agent further conducted analyses aiming to map the OSMR-associated transcriptional program, finding evidence for receptor amplification, extracellular matrix remodeling, and immune cell recruiting chemokines in the gut as a fibroblast’s OSMR expression increased (Figure 5D). These insights help inform how OSMR signaling in certain fibroblast states contribute to continued inflammation in UC non-responders, suggesting a stromal signaling environment that could be less dependent on TNF-driven inflammation.

Synthesizing these findings, the virtual CSO hypothesized that the futility of the MOONGLOW trial may reflect a precision medicine gap: vixarelimab was evaluated in an unselected UC population without biomarker-based enrichment for OSMR expression, even though the trial’s own protocol recognized that elevated OSMR is associated with poor response to anti-TNF therapy. Notably, in current clinical practice, UC treatment selection and sequencing are still driven primarily by disease severity and prior treatment exposure within a rapidly expanding, approval-driven therapeutic portfolio, rather than by validated prospective molecular stratification [71–78]. Consequently, dose escalation and switching across therapeutic mechanisms are common until an effective option is achieved. To date, no published studies have explicitly proposed OSMR as a stratification biomarker for selecting patients for anti-OSMR clinical trials in UC. Based on these data-driven insights, the virtual CSO proposed revising the trial design to use biomarker-guided enrollment, thereby enriching the study population with patients exhibiting high OSMR expression, which vixarelimab may be particularly effective in targeting (Figure 5E).

To assess feasibility, the virtual CSO consulted the clinical trialist agent, and it reasoned that such biomarker stratification appears feasible in practice: UC mucosal biopsies are already routine during standard-of-care colonoscopy, and the infrastructure for immunohistochemistry or gene expression is mature. Additionally, the agent observed that the current primary endpoint (Week 12 clinical remission by modified Mayo Score) is a coarser clinical composite that may be insensitive to OSMR pathway engagement, which could obscure a genuine responder signal in an unselected population and when assessed at a single futility timepoint. Incorporating a prespecified biomarker-based target engagement endpoint to complement clinical or endoscopic outcomes could therefore enhance interpretability. The clinical trialist agent provided additional justification by referring to precedent of biomarker enrichment in oncology trials such as PD-L1 selection in KEYNOTE-024 (NSCLC), HER2 testing for trastuzumab (breast cancer), and EGFR screening (NSCLC) [79–81]. Finally, it flagged some design decisions for a potential biomarker-enriched OSMR trial which included defining clinically meaningful OSMR expression cutoff levels, estimating the prevalence of OSMR-high patients to anticipate screen-failure rates, and selecting the appropriate assay platform.

## Discussion

The Virtual Biotech is a multi-agent platform for early-stage drug discovery. At its core, a virtual CSO plans and coordinates multiple domain-specific scientist agents that analyze and reason over primary data sources. The domain-specialized scientist agents mirror core R&D functions and have access to a wide variety of tools and data sources. This collaborative setup allows for unified decision-making based on a collective and nuanced understanding of many primary data sources. Our experiments show that the Virtual Biotech is capable of performing large-scale analysis of clinical trials, and identified several novel transcriptomic features of drug targets associated with clinical trial success. Additionally, two case studies on real-world drug development scenarios show that the Virtual Biotech can yield insightful hypotheses that complement the evidence and conclusions reached by modern biopharmaceutical companies.

Compared to a traditional manual target validation workflow, the Virtual Biotech offers three main benefits. First, because traditional workflows are often siloed, teams often work independently with little reference to data generated by other teams. This approach can miss critical opportunities. For example, in the B7-H3 lung cancer case study, a reviewer agent’s critique that ligand–receptor inferences from dissociated single cell data lacked spatial context triggered a follow-up spatial transcriptomics analysis to test whether B7-H3 regions correspond to immune exclusion. The virtual CSO then handed off to the clinical trialist agent to quantify clinical relevance via biomarker-stratified survival modeling in TCGA LUAD. In siloed workflows, these cross-division handoffs may not occur and therefore may produce incoherent conclusions. The Virtual Biotech’s ability to deeply integrate these diverse data sources early in the drug discovery process promises to enable better informed decision-making before drugs are taken to clinical trials.

Second, the Virtual Biotech has the potential to support more objective, data-driven decisionmaking than is possible in large human teams. In biopharmaceutical companies, reaching consensus can be challenging, as organizational dynamics, differing prior beliefs, and other non-technical considerations may sometimes shape judgment alongside the evidence. The objective and datadriven nature of the Virtual Biotech means it can make recommendations unbiased by sociological influences. Additionally, the transparent documentation of Virtual Biotech-driven analyses makes it easy for human experts to audit its findings, helping support more reproducible research compared to the often fragmented or lost audit trails in manual workflows.

Third, the speed and scalability of the Virtual Biotech is much greater than traditional workflows. The B7-H3 analysis involved drawing from genetics, transcriptomics, and clinical data to reach a nuanced assessment of target disease biology and therapeutic modality, but took less than one day and cost $46.00 in API fees. Similarly, the in-depth analysis of a terminated trial testing OSMR*β* as a target in UC took less than one day, costing $54.00 in Anthropic API credits. This efficiency in terms of both time and cost permits scaling of analyses much more than otherwise possible. For example, manual annotation of clinical trial outcomes would be prohibitive, but the Virtual Biotech’s large-scale analysis revealed novel insights that are only possible at that scale. Along these lines, the Virtual Biotech could enable deep analysis of many potential targets for a given disease area at the level of rigor showcased in the case studies.

The single-cell features of drug targets identified by the Virtual Biotech are themselves of independent interest as new predictors of clinical trial success. Genetic evidence has long been recognized as one of the strongest predictors of clinical success and now plays a central role in modern target prioritization [33]. Prior work has also suggested that single-cell–derived signals contain meaningful clinical information, linking these features to clinical phase progression [47]. Here, we build on these observations by demonstrating that single-cell–based features are not only associated with phase advancement but also with successful endpoint outcomes and reduced adverse event rates across multiple organ systems. This extension is important, as endpoint success provides a more direct and clinically meaningful measure of therapeutic efficacy rather than a surrogate marker of development progress. Furthermore, we showed that single-cell evidence is orthogonal to genetic evidence and that significant associations were replicated within trials with no supporting genetic evidence. Expression-based features can be especially valuable in oncology because they better capture tumor state and microenvironment biology that is not explained by germline variation alone. Together, these findings suggest that incorporating single-cell–derived evidence alongside genetic support may provide important guidance in drug development programs.

Our current study has limitations. AI agents can make analysis or interpretation errors [13]. As in traditional drug discovery settings, a single mistake can propagate through the virtual organization. Although we introduce a scientific reviewer agent to mitigate this concern, it is still pertinent that human experts are involved in the workflow. We note that one benefit of the Virtual Biotech is that it allows human scientists to precisely audit the agent-generated unified end-to-end workflow and reasoning. This level of transparency is rarely achievable in a traditional drug research environment, where analytical or interpretive errors can propagate without a clear or accessible audit trail.

Additionally, our trial analyses are observational and should be interpreted as such. Possible confounders include trial targets, indications, endpoint definitions, and sponsor-related influences. In this vein, we adjusted for a trial’s phase, year of enrollment, therapeutic area, and drug modality to address their potential confounding, observing consistent results. In addition, the analysis was limited to trials registered in the ClinicalTrials.gov database and drugs which have mapped molecular targets. For the case studies, the Virtual Biotech generated data-driven hypotheses, which must be tested and further validated. There is the possibility of information leakage due to pretraining of the language models, biasing the agent towards existing hypotheses. To mitigate this, we intentionally selected case studies with trial readouts that occurred months after the language model knowledge cutoff date of January 2025. We also verified that the specific analysis and findings from the agents have not been previously reported in literature.

In the future, we envision expanding the Virtual Biotech to encompass even more of the drug discovery pipeline. This could include capabilities such as molecule design, virtual screening, lead optimization, toxicity prediction, and even clinical decision-making agents, bringing the benefits of interdisciplinary collaboration, speed, and objectivity to the entire end-to-end process. The Virtual Biotech illustrates a shift from isolated AI tools toward coordinated systems that reason across biological scales and stages of translation. As agentic platforms mature, they may enable a new model of scientific organization in which hypotheses are continuously generated, stress-tested against heterogeneous evidence, and refined in partnership with human experts. Rather than replacing scientists, such systems could expand the scope and speed of therapeutic exploration while making the reasoning process more transparent and reproducible.

## Data Availability

All data used in this study are publicly available. Data from the Open Targets Platform can be directly accessed at https://platform.opentargets.org. CELLxGENE data can be accessed at https://cellxgene.cziscience.com. The Tabula Sapiens dataset can be accessed at https://tabula-sapiens.sf.czbiohub.org/. The Visium LUAD dataset can be accessed at https://pubmed.ncbi.nlm.nih.gov/38782901/. The TCGA PanCancer Atlas LUAD cohort can be accessed at https://www.cbioportal.org/. The microarray data for the five UC trials analyzed can be access with GEO accession IDs GSE12251, GSE16879, GSE92415, GSE206285, and GSE73661. The TAURUS dataset can be accessed at https://pubmed.ncbi.nlm.nih.gov/39438660/.

## Acknowledgments

The authors thank the members of the Zou lab at Stanford University and Wei Deng for their input and useful discussions on this work. H.G.Z. is supported by the Knight-Hennessy Scholarship and the Stanford University Medical Scientist Training Program (NIH T32-GM145402). P.E. is supported by the Stanford Graduate Fellowship and the NSF GRFP (NSF DGE-2146755). J.Z. is supported by the Chan Zuckerberg Biohub.

## Extended Methods

### Virtual Biotech System Architecture

Virtual Biotech is a multi-agent AI system that is organized to mimic the structure of a biotechnology research and development organization to accelerate and support therapeutic development and strategy. The system employs a hierarchical orchestration model comprising a Chief Scientific Officer (CSO) as the global strategic orchestrator who oversees domain-specialized AI scientist agents and dedicated research support agents.

The virtual CSO agent supports multi-turn conversations with users, interprets queries, and dispatches analytical tasks to appropriate scientist agents. The CSO functions exclusively as an orchestrator: it routes queries, synthesizes findings, and provides strategic recommendations, but never directly accesses data or performs analyses. This separation of roles ensures that domain expertise remains encapsulated within scientist agents while the CSO maintains a global view of the research process.

### Human Interaction with the Virtual Biotech

Virtual Biotech is designed for lightweight human interaction, in which the primary role of the user is to set intent and guide the system through high-level scientific questions. After a human user submits a high-level scientific query, the CSO agents conducts a brief clarification interview— posing follow-up questions to align on scope feasibility—before initiating computational analyses (Figure 1C). After objectives are clarified, subsequent actions are autonomous: the CSO decomposes the task and routes sub-tasks to scientist agents, scientist agents perform analyses and reason over findings, and the CSO synthesizes cross-division results into recommendations. Consistent with this design, in the case studies presented in this paper, human involvement was limited to (i) responding to CSO clarification prompts and (ii) asking follow-up questions after each iteration (e.g., to probe assumptions, request sensitivity checks, or trigger additional analyses). Experimental ideation, design, and execution remained agent-driven. Users can independently verify key results by inspecting and executing the generated code, enabling human oversight.

### AI Scientist Agents

The Virtual Biotech consists of eight distinct scientist agents distributed across four research divisions, plus three additional agents in the CSO’s office that oversee the scientific direction and quality of the work (Figure 1A). Each agent is configured with domain-specific system prompts and MCP access.

- **Target Identification and Prioritization Division**: The statistical genetics agent evaluates human genetic evidence by querying genome-wide association studies (GWAS), locus-to-gene (L2G) machine learning predictions for causal gene assignment, fine-mapped credible sets, rare variant burden meta-analyses, and quantitative trait locus (QTL) colocalization to determine whether genetic variation affecting a gene influences disease risk. The functional genomics and perturbation agent assesses target essentiality and pharmacological tractability using CRISPR knockout screens and drug perturbation transcriptomic profiles from the Tahoe-100M dataset to evaluate whether a target can be modulated. The single cell atlas agent queries CELLxGENE Census, which contains 143 million harmonized single cell profiles, to learn cell-type-specific expression patterns, differential expression between disease and healthy states, and cell-cell communication patterns
- **Target Safety Division**: The bio-pathways and protein-protein interaction agent reasons through the target’s biological context through pathway and molecular interaction maps to identify involvement in critical cellular processes that may pose safety concerns upon modulation. The single cell atlas agent complements this network-level assessment by examining cell-type-specific expression patterns across cell types found in 27 tissues of the Tabula Sapiens v2 atlas to identify potential off-target toxicity liabilities in critical cell types [40]. The FDA safety officer agent complements these mechanistic assessments with empirical safety data from OpenFDA adverse event reports for drugs with similar mechanisms, drug safety warnings and black box labels from drug labels, and mouse knockout phenotypes to reason through potential target-level safety liabilities.
- **Modality Selection Division**: The target biologist agent evaluates target properties that constrain modality choice, including protein family classification, subcellular localization, and structural tractability predictions that assess the likelihood of developing small molecule binders, therapeutic antibodies, or other modality types based on protein structure and binding site accessibility. The pharmacologist integrates this biological evaluation with existing clinical precedence, querying ChEMBL-derived drug databases to find approved therapies, pipeline compounds, and chemical probes that share the same target or mechanism. In addition, they analyze homologous proteins to determine family-level tractability and assess development practicality for each modality, including projected timelines, manufacturing demands, and likely hurdles.
- **Clinical Officers Division**: Working in tandem with the FDA safety officer agent, the clinical trialist agent enhances regulatory assessment with clinical precedence analysis. It locates prior trials aimed at the same gene or pathway, extracts information on trial phase, enrollment, endpoints, and results to determine the level of clinical validation, and examines discontinued studies to identify reasons for failure, including futility, safety concerns, or business-related decisions. For oncology targets, the clinical trialist agent can access clinicogenomics data to determine prognostic biomarkers across tumor types and survival outcomes from cBioPortal.
- **Office of CSO Agents**: The Chief of Staff provides rapid strategic intelligence briefings on research fields and data landscape by inventorying MCP tools and performing web searches. The scientific reviewer performs quality assurance on AI scientist agent outputs, identifying gaps, unsupported conclusions, or misalignment with user queries (Supplementary Note II).

All agents operate using Claude Sonnet 4.5, except for the Chief of Staff and Scientific Reviewer, which use Claude Haiku 4.5 for efficiency and speed. Scientist agents have access to standard computational tools including file operations and code execution, plus domain-specific data access tools in the form of MCP servers (described below). Scientist agents are instructed to maintain critical scientific thinking by questioning assumptions, validating findings across complementary datasets, and classifying evidence strength as weak or strong based on the convergence of independent lines of evidence.

### Implementation and Language Model Configuration

Agents and orchestration were implemented using the Claude Agent SDK, an Anthropic agent development framework to build agent AI platforms [82]. The SDK provides several capabilities essential for multi-agent scientific workflows: hierarchical agent orchestration enabling a parent agent to spawn and coordinate child agents with independent execution contexts, native integration with the MCP for standardized tool access, and built-in support for multi-turn conversations with persistent conversation state. These capabilities allow the Virtual Biotech to implement complex delegation patterns where the CSO orchestrator manages scientist agents that operate autonomously while maintaining coordinated access to shared data resources. Agent behavior is controlled through structured system prompts that define role identity and expertise boundaries, available tools and data access permissions, analysis workflows and decision frameworks, and output formatting requirements. Specialized scientific workflows, such as single cell qualitycontrol and analysis, were implemented in the form of agent Skills with progressive disclosure of workflow guidance.

### Data and Virtual Biotech Tool Infrastructure

The Virtual Biotech integrates data from multiple public biomedical databases to enable a comprehensive evaluation of drug targets across the translational scientific spectrum and multiple biological scales. The Open Targets Platform (version 25.09) serves as the primary data backbone [27, 36]. This is an extensive database that enables systematic discovery and evaluation of candidate therapeutic drug targets. By combining publicly available datasets, including those produced by the Open Targets experimental and informatics research programs, Open Targets curates data that support the development of therapeutic hypotheses. Each component contributes specific evidence types and detailed, high-resolution information on drug targets, diseases, genetic variants, QTL colocalization, mechanisms of action of therapeutic compounds, scientific publications, drug labels, FDA-reported safety signals, clinical trials, and other related data [27, 36].

We further integrated additional data resources into the Virtual Biotech that are key evidence sources in drug discovery. Single-cell atlas data are accessed and retrieved through MCP tools that leverage the CELLxGENE Census API, which aggregates harmonized single-cell RNA-sequencing profiles from over a thousand published studies, offering standardized cell type annotations across diverse tissues and disease conditions [50]. In addition, we provided the Virtual Biotech with a local copy of the Tabula Sapiens v2 healthy reference atlas [40]. Pseudobulked differential expression profiles were procured from Tahoe-100M, a giga-scale single-cell perturbation atlas of 50 cancer lines exposed to 1,100 small molecule perturbations [28]. Agent access to clinicogenomic and outcome data in different cancer cohorts was implemented using MCP tools powered by the cBioPortal REST API [83]. Finally, clinical trial information was made available to agents via tools that leveraged the ClinicalTrials.gov API [84].

Importantly, data access to each agent is mediated through ten MCP servers, each implemented as a FastMCP Python instance that exposes domain-specific query functions as callable tools (Figure 1B). MCP provides a standardized interface between language models and external data sources, enabling structured queries with typed parameters and formatted responses [23, 24]. Each MCP tool is implemented as a Python function with comprehensive docstrings and typed parameter signatures, which FastMCP automatically introspects to generate structured tool schemas exposed to the language model at runtime, injecting required context that enables agents to discover available tools and understand their purpose, required inputs, and expected outputs without manual specification [85]. Tools provide not only raw outputs but also interpretable summaries and preview results, allowing agents to rapidly gauge relevance and determine whether more in-depth analysis is warranted. This facilitates efficient multi-step reasoning and iterative tool invocation across diverse data sources. Each MCP server offers specialized tools tailored to frequent analytical tasks. These tools accept structured inputs—such as gene symbols, disease identifiers, and various filter parameters—and return well-formatted outputs optimized for language model consumption. The Virtual Biotech MCP tools are intentionally composable, enabling agents to chain multiple tool invocations together to construct rich, comprehensive evidence profiles.

### Benefits of multi-agent architecture

In contrast to single-agent approaches, the multi-agent architecture of the Virtual Biotech offers several key advantages. The breadth of data sources and analytical tools incorporated into the Virtual Biotech, on the scale of hundreds of tools and terabytes of data spanning ∼80,000 targets, 40 million papers, and 100 million cell profiles, makes in-depth analyses of these sources by a single agent infeasible due to context, time, and memory constraints [27, 28, 86]. By contrast, a multi-agent framework allows these analyses to be decomposed and distributed across parallel agents with isolated contexts, enabling each agent to conduct a thorough, focused analysis without overwhelming the shared context. High-level summaries from these agents are then integrated by a coordinating agent to support final decision-making.

Second, the multi-agent architecture enables parallel workflows at scale. For example, in extracting clinical trial outcomes, each extraction followed a multi-step workflow: given an input NCT ID, the clinical trialist agent first calls the ClinicalTrials.gov API to retrieve trial metadata, then evaluates whether all required fields are sufficiently populated before moving through a three-tier evidence cascade. This cascade involves searching PubMed with NCT ID confirmation and then expanding to additional web sources. Throughout this process, the agent reviews numerous information sources to understand trial designs, interpret complex clinical outcomes, determine endpoint results, map termination reasons to a standardized ontology, and consolidate all evidence into a validated JSON output that adheres to a Pydantic schema. Consequently, we assigned a dedicated agent to each trial, allowing it to use its full context window on that single study—gaining deep familiarity with its design, results, and literature—without the distraction or context dilution of managing multiple trials in one conversation. Additionally, this design enables our multi-agent parallelized workflow where 37,075 phase II and phase III trials were systematically processed in 6 hours. In contrast, processing the same number of trials with a single agent would require approximately 1,839 hours or 76.6 days of runtime. The parallelized multi-agent architecture therefore achieves an approximately 184-fold speedup, reducing a months-long extraction to an overnight run.

### Analysis of Single-Cell Features and Clinical Trial Outcomes

#### Clinical trial manual validation

We sampled 100 trials from the Open Targets clinical trial dataset that was included in the agent-driven curation process (50 randomly sampled from all Phase II trials, and 50 from all Phase III). Two human annotators assessed agreement with the agent’s annotations for primary outcomes, secondary outcomes, and adverse event (AE) rates. For each trial, the annotators first reviewed the ClinicalTrials.gov record for posted results and any linked publications; if none were available, they searched Google by NCT identifier and trial name to identify relevant abstracts or reports. The annotators then compared these sources against the agent’s extracted outcomes, performing additional review when discrepancies arose and documenting the reasons for disagreement. The two human annotators discussed ambiguous cases to reach a consensus.

#### Single-Cell Feature Extraction from Tabula Sapiens

Single-cell expression features were derived from the Tabula Sapiens atlas, comprising 27 human tissues (Bladder, Blood, Bone Marrow, Eye, Fat, Heart, Kidney, Large Intestine, Liver, Lung, Lymph Node, Mammary, Muscle, Pancreas, Prostate, Salivary Gland, Skin, Small Intestine, Spleen, Stomach, Testis, Thymus, Tongue, Trachea, Uterus, Vasculature, and Ear). Cell type annotations were based on the Cell Ontology class assignments provided in the atlas. For each target gene, features were first computed per tissue and then aggregated across tissues to obtain a single global value per gene.

#### Cell Type Specificity: Tau Index

Cell type specificity was quantified using the Tau index (*τ*), which measures the degree to which gene expression is restricted to specific cell types [41]. For a gene with mean expression *x̄_j_*in cell type *j* across *n* cell types within a given tissue:

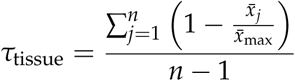

where *x̄*_max_ = max*_j_*(*x̄_j_*) is the maximum mean expression across all cell types within that tissue. The Tau index ranges from 0 (ubiquitous expression, equal across all cell types) to 1 (perfectly specific expression, expressed in only one cell type). Cell types with fewer than 20 cells were excluded from Tau computation, and mean expression was computed from log-normalized counts. For a given gene target, the Virtual Biotech then aggregated Tau metrics across all tissues as the arithmetic mean of per-tissue Tau values across tissues in which the gene was expressed.

#### Expression Heterogeneity: Bimodality Coefficient

Within-population expression heterogeneity was assessed using the bimodality coefficient (BC), which detects the presence of distinct expression subpopulations [42, 43]. For a gene’s expression vector across *n* expressing cells (cells with expression *>* 0) within a given tissue:

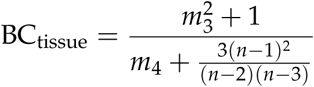

where *m*_3_ is the sample skewness and *m*_4_ is the Pearson kurtosis. The bimodality coefficient ranges from 0 to 1, with BC *>* 0.555 suggesting a bimodal distribution. Only expressing cells were included in the computation to focus on heterogeneity among cells that transcribe the gene. The global bimodality score for each gene was computed as the arithmetic mean of per-tissue bimodality coefficients.

#### Tahoe-100M Hallmark Signature Scores

Drug-induced transcriptional responses were quantified using six hallmark signature scores derived from the Tahoe-100M perturbation atlas. For each drug-cell line combination, log_2_ foldchange (LFC) values were extracted at maximum dose, and non-significant changes (adjusted *p* ≥ 0.05) were set to zero. Hallmark scores were computed as the mean LFC across curated gene sets, with sign adjustments to ensure positive scores indicate drug efficacy:

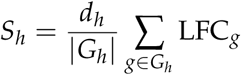

where *G_h_* is the gene set for hallmark *h* and *d_h_* ∈ {−1, +1} is a direction coefficient. The six hallmarks were: (1) *Apoptosis* (*d* = +1; 11 pro-apoptotic genes including BAX, CASP3, CASP9), (2) *Proliferation suppression* (*d* = −1; 11 proliferation markers including MKI67, PCNA, TOP2A), (3) *DNA damage response* (*d* = +1; 5 p53 targets including GADD45A, MDM2), (4) *Stress response* (*d* = +1; 11 ER/oxidative stress markers including DDIT3, ATF4, HSPA5), (5) *Resistance* (*d* = +1; 8 anti-apoptotic genes including BCL2, MCL1, XIAP), and (6) *Cell cycle arrest* (CDKN1A, CDKN1B, CDKN2A, BTG2, *d* = +1; and CCNA2, CCNB1, CCNE1, *d* = −1). These gene sets were directly selected by the functional genomics agent when designing its analysis based off the each gene’s canonical roles in these pathways. For all scores except resistance, positive values indicate the expected direction of drug efficacy; for resistance, positive values indicate potential resistance mechanisms.

#### Statistical analysis of single cell and perturbation atlas features with trial outcomes

Our statistical analyses of single cell features followed the same approach in analogous studies of genetic evidence [33, 34]. Clinical trial outcomes (phase progression, trial termination, endpoint success) were analyzed with univariate logistic regression to estimate the association between single-cell expression features and outcome probability. Features were z-score standardized prior to model fitting to enable comparison of effect sizes across continuous features with different scales. Model parameters were estimated via maximum likelihood using generalized linear models with a binomial family and logit link (statsmodels.GLM, Binomial family). For trials assessing drugs that act on multiple molecular targets, and thus yield several tau and bimodality features per trial, the minimum value across all targets for each feature type was used as the single cell agent hypothesized that the least specific target likely drives the overall safety profile. For continuous percentage outcomes such adverse event rates by organ system calculated at the patient-level, we used beta regression as implemented in R’s betareg package [87]. The resulting regression coefficient corresponds to the change in the logit of the mean adverse event proportion associated with a one-standard-deviation increase in the predictor. These univariate analyses are the main results we present, consistent with the approaches of Razuvayevskaya et al. and Minikel et al [33, 34].

In addition to these results, we conducted robustness analyses to further support our findings. First, to assess whether observed associations exceeded chance, we conducted permutation tests with 1,000 iterations per feature-outcome combination. In each iteration, the outcome vector was randomly shuffled while retaining the original feature values, and the regression was re-fit to obtain a null coefficient. Empirical two-sided *p*-values were computed as the proportion of null coefficients with absolute value greater than or equal to the observed coefficient. Permutation tests were run for both the logistic (binary outcome) and beta regression (adverse event rate) analyses using identical model specifications as the primary analyses.

Next, to account for potential confounding by drug modality, therapeutic area, trial start year, and clinical phase, we fit generalized mixed-effects models (GMMs). For binary outcomes, we used logistic GMMs via lme4::glmer in R with crossed random intercepts for drug-type combination and therapeutic area combination. For adverse event rate outcomes, we used beta mixedeffects regression via glmmTMB::glmmTMB with a beta family and the same crossed random-effects structure.

To evaluate whether single-cell expression features provided additional information independent of genetic evidence, we fit bivariate logistic regression models adjusting for genetic evidence. For adverse event outcomes, the analogous bivariate beta regression was fit using R’s betareg. As a complementary analysis, we repeated the univariate analyses restricting to the subset of 42,165 trials (74.6%) that have no genetic evidence. Following the approach of Razuvayevskaya et al., genetic evidence was defined as a binary indicator of whether any target–disease indication pair in a trial had a direct genetic association reported in Open Targets, which aggregated evidence from genome-wide association studies, phenome-wide association studies, gene burden analyses, and clinically annotated variants (ClinVar/EVA) [27, 34].

To adjust for multiple hypothesis testing, we used the Benjamini-Hochberg procedure to control for a false discovery rate of 5% [88].

To enable interpretable quantification, we also binarized the continuous tau cell-type specificity index into either "cell-type-specific" or "broadly expressed" categories using K-means clustering (*k* = 2) on the trial-level distribution, setting the threshold at the midpoint of the two cluster centers (*τ* = 0.69).

### Single-Cell and Spatial Transcriptomics Analyses

#### Quality Control and Data Preprocessing

Raw single-cell or single-nucleus RNA-sequencing count matrices were subjected to quality control filtering. Cells were retained if they expressed between 300 and 9,000 genes and had ≤15% mitochondrial reads. To mitigate compositional biases from uneven sampling, donor-level downsampling was performed (maximum 10,000 cells per donor), stratified by cell type to preserve relative proportions. Gene expression was normalized using library-size normalization and log1p transformation via Scanpy (v1.9+) [89]. Highly variable genes (3,000) were selected for downstream dimensionality reduction.

Cell type annotations across datasets were harmonized to a common vocabulary using the Cell Ontology [90]. Fine-grained annotations were mapped to standardized Level 3 ontology categories through hierarchical ontology traversal. Cell types were filtered to those eligible for downstream differential expression analysis, requiring representation by ≥3 donors per condition and ≥20 cells per donor.

Dimensionality reduction was performed using principal component analysis (50 components). Batch effects were corrected using Harmony [91] to preserve biological variation while mitigating technical differences across datasets. Uniform Manifold Approximation and Projection (UMAP) embeddings were computed on the Harmony-corrected principal components for visualization. Batch mixing was assessed by comparing corrected and uncorrected embeddings.

#### Pseudobulk Differential Expression

Differential expression analysis was performed using a pseudobulk approach. Raw counts were aggregated by donor and cell type, and differential expression was tested using PyDESeq2 [92]. Genes were considered differentially expressed at FDR *<* 0.05 and |log_2_FC| *>* 0.5.

#### Cell-Cell Communication Analysis

Cell-cell communication inference was performed using LIANA [51], which computes a consensus score across multiple ligand-receptor inference methods. LIANA’s rank_aggregate method was run with 1,000 permutations for each group. Interactions were retained if they ranked in the top 10%, had *p <* 0.01, and were supported by ≥3 constituent methods. Group-specific interactions were identified by comparing interaction sets that met the aforementioned criteria between conditions.

#### Cell Type Deconvolution of Spatial Transcriptomics Data

Cell type deconvolution of 10x Visium spatial transcriptomics data was performed using Cell2Location on a NVIDIA H100 GPU [53]. Briefly, the workflow first trained a negative binomial regression model on the LUAD single-cell atlas to estimate gene expression signatures for each cell type. The trained reference signatures were then applied to spatial data to infer cell type abundances per spot, with an expected cell density of 8 cells per spot. We excluded the gene of interest (B7-H3) from this workflow to avoid circularity in the downstream analyses. The reference model was trained for 250 epochs and the spatial mapping model for 10,000 epochs. Cell type proportions were computed from the top fifth percentile posterior estimates of cell abundances.

#### Spatial Immune Neighborhood Analysis

To assess immune cell composition in the local microenvironment of gene-expressing spots, a *k*-nearest neighbor graph (*k* = 6) was constructed for each tissue section [93]. Within each sample, expressing spots were stratified into gene-high (top quartile) and gene-low (bottom quartile) groups based on per-sample thresholds. Samples with fewer than 25 expressing spots were omitted. For each spot, the mean Cell2Location-estimated cell-type abundance across its *k* nearest spatial neighbors was computed. To test for immune depletion while controlling for confounders of spatial tissue architecture, mixed-effects models were fit separately for each immune cell type:

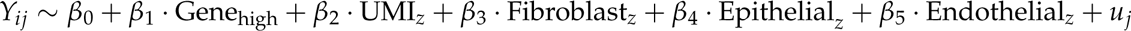

where *Y_ij_* is the mean neighbor immune abundance for spot *i* in sample *j*, total UMI is as a proxy for sequencing quality, and fibroblast, epithelial, and endothelial cell abundances control for spatial compartment, and *u_j_* ∼ N (0, *σ*_*u*_^2^) is a sample-level random intercept to account for slide-to-slide variation. Models were estimated using restricted maximum likelihood (REML) via statsmodels [94].

#### Pathway Activity Inference and Mixed-Effects Modeling in Ulcerative Colitis (UC)

To assess signaling pathway dynamics in UC tissue before and after biologic treatment, pathway activity scores were inferred from the TAURUS longitudinal single-cell atlas of anti-TNF (adalimumab) treatment in inflammatory bowel disease [70]. The analysis focused on UC patients (n=22) with paired pre- and post-treatment colonic biopsies. Raw counts were normalized (library-size normalization to 10,000 counts per cell, log1p transformation) and pseudobulked by computing the mean expression for each combination of cell state and sample, retaining only combinations with ≥10 cells. PROGENy pathway activity scores were then calculated for 14 canonical signaling pathways using the multivariate linear model method implemented in the decoupler Python package with the top 100 pathway-responsive genes per pathway from the human PROGENy model [95].

To evaluate whether changes in pathway activity following treatment differed between responders and non-responders, linear mixed-effects models were fit in R using lme4::lmer. Treatment (Pre/Post) and remission status (Remission/Non-Remission) were included as fixed effects along with their interaction, site (biopsy location) was modeled as an additional fixed effect, and each patient was modeled with random effects to account for repeated measurements within individuals.

#### OSMR Expression Gradient Analysis

To characterize the transcriptional program associated with increasing OSMR expression in UC fibroblasts, an OSMR expression gradient analysis was performed on the three fibroblast cell states in which OSMR was significantly upregulated in non-responders after anti-TNF treatment as previously reported [70]. Cells from UC non-responder patients in these fibroblast states were pooled, and each cell was assigned an OSMR gradient value by rank-normalizing its OSMR expression to the [0, 1] interval. For each gene that was expressed in at least 10% of cells, we fit a linear mixedeffects model using lme4::lmer. In this model, each gene’s expression was the outcome, and OSMR_rank (rank-normalized OSMR expression) was the primary predictor. Sample treatment status (pre/post), age (z-scored), gender, biopsy site, and the number of detected genes (z-scored, used as a proxy for cell quality) were included as fixed-effect covariates, while patient identity was incorporated as a random effect. P-values for the OSMR_rank coefficient were corrected for multiple testing using the Benjamini–Hochberg procedure (FDR *<* 0.05). [88].

### Biomarker-Stratified Survival Analysis

To assess the prognostic significance of B7-H3 expression in LUAD, survival analysis was performed on the TCGA PanCancer Atlas LUAD cohort accessed via cBioPortal [54, 55]. Gene expression profiles and clinical annotations—including age at diagnosis, AJCC pathologic tumor stage, sex, and survival endpoints—were retrieved for 566 LUAD patients. Analysis was restricted to patients with complete covariate data for age, stage, and sex. Patients were stratified into expression quartiles based on B7-H3 (CD276) expression, comparing the top quartile (high expression) against the bottom quartile (low expression). Multivariable Cox proportional hazards models were fit for overall survival (OS), progression-free survival (PFS), disease-specific survival (DSS), and disease-free survival (DFS), adjusting for age (continuous), cancer stage (advanced [III/IV] vs. early [I/II]), and sex. Survival analyses were performed using the lifelines Python package.

### Cross-Trial Analysis of Baseline OSMR and OSM Expression

To evaluate whether baseline mucosal OSMR and OSM expression levels are associated with biologic treatment response in UC, gene expression data from five independent clinical trial cohorts were analyzed. For each dataset, pre-processed expression matrices and sample-level metadata were obtained directly from the Gene Expression Omnibus (GEO). When multiple microarray probes mapped to the same gene, expression was summarized as the arithmetic mean across all available probes for that gene. Patients were classified as responders or non-responders according to each trial’s protocol-defined clinical and endoscopic endpoints. Baseline expression levels were compared between responders and non-responders using the two-sided Mann–Whitney test.

## Supplemental Figures

**Figure S1:**
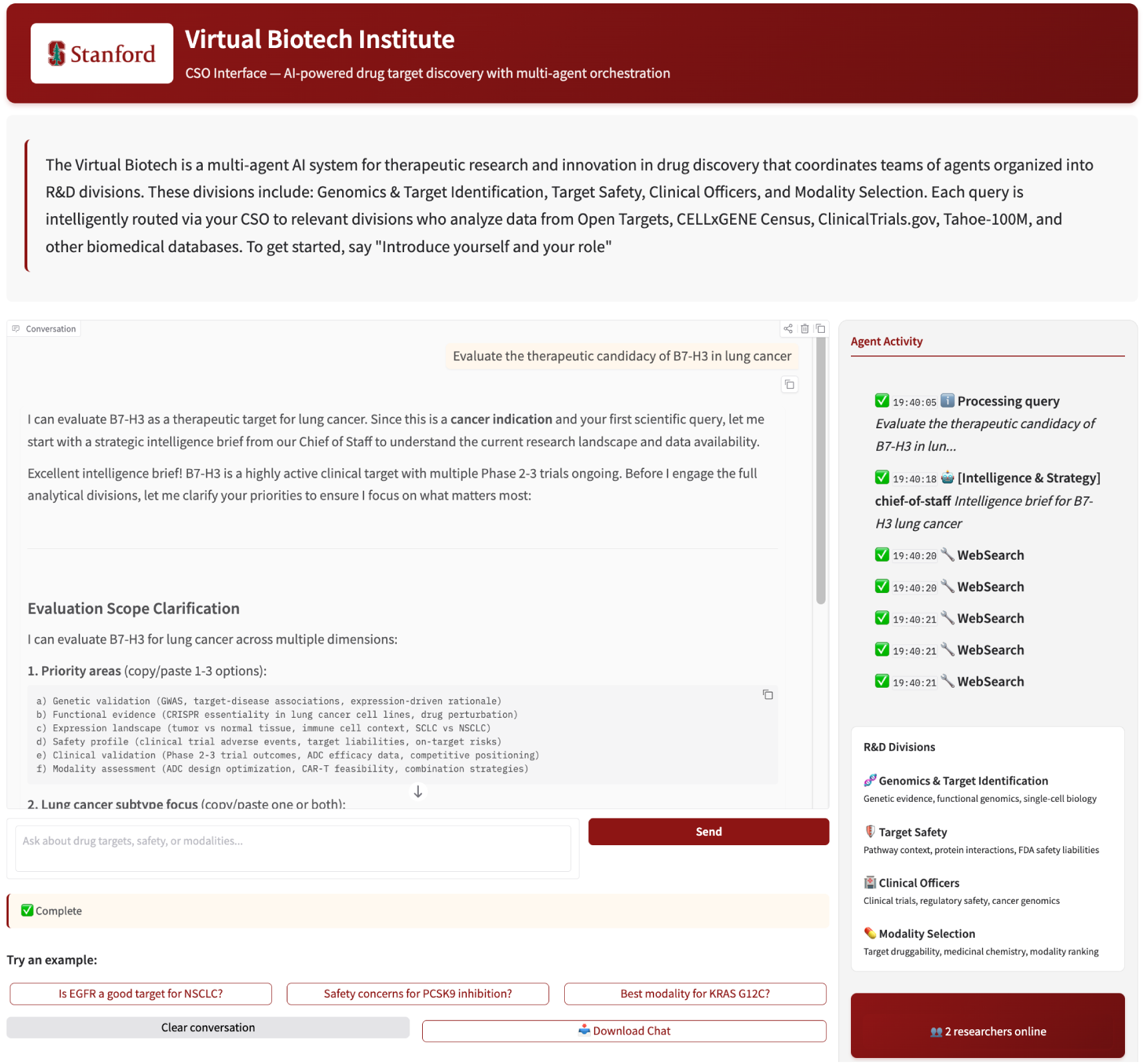
Virtual Biotech User interface. The user interface enables natural language queries about drug targets, safety, and therapeutic modalities through a chat-based system powered by a Chief Scientific Officer agent that coordinates AI scientist agents. Real-time activity tracking displays which agents are analyzing data from biomedical databases and their reasoning traces, with all generated code, figures, and data files accessible for download to facilitate reproducibility. Each user session maintains an isolated workspace for reproducible analysis and file management.

**Figure S2:**
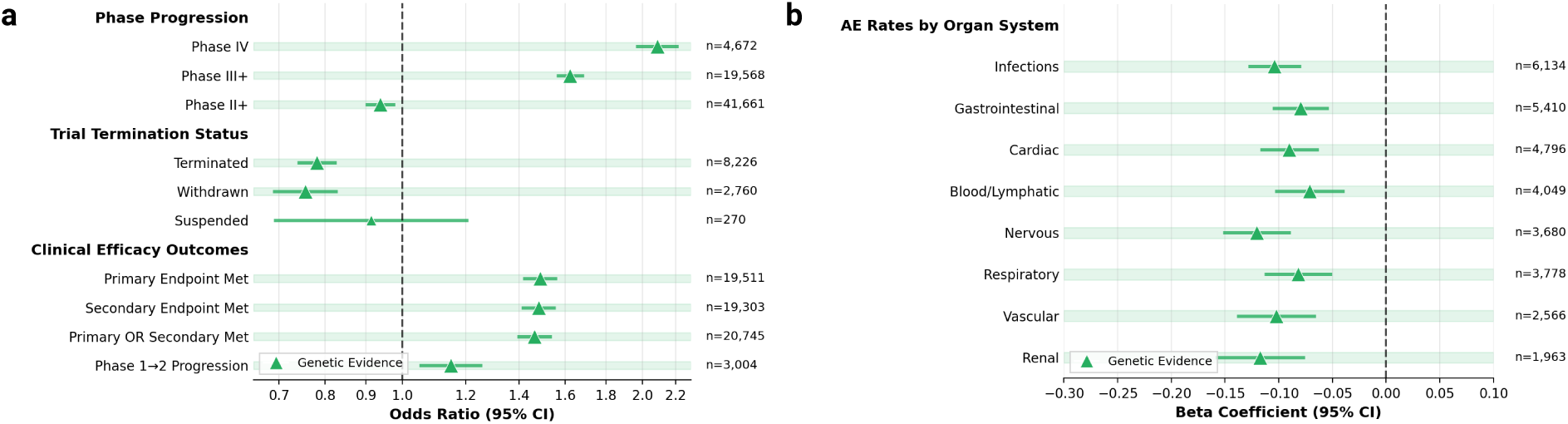
Genetic evidence of drug targets from Open Targets platform and their associations with clinical trial outcomes. (A) Associations between genetic evidence and phase progression, trial stoppage, and efficacy outcomes. Odds ratios (ORs) estimated with logistic regression. Error bars represent 95% confidence intervals. (B) Associations between genetic evidence and serious adverse event rates grouped by organ system. Beta coefficients estimated with beta regression. Error bars represent 95% confidence intervals.

**Figure S3:**
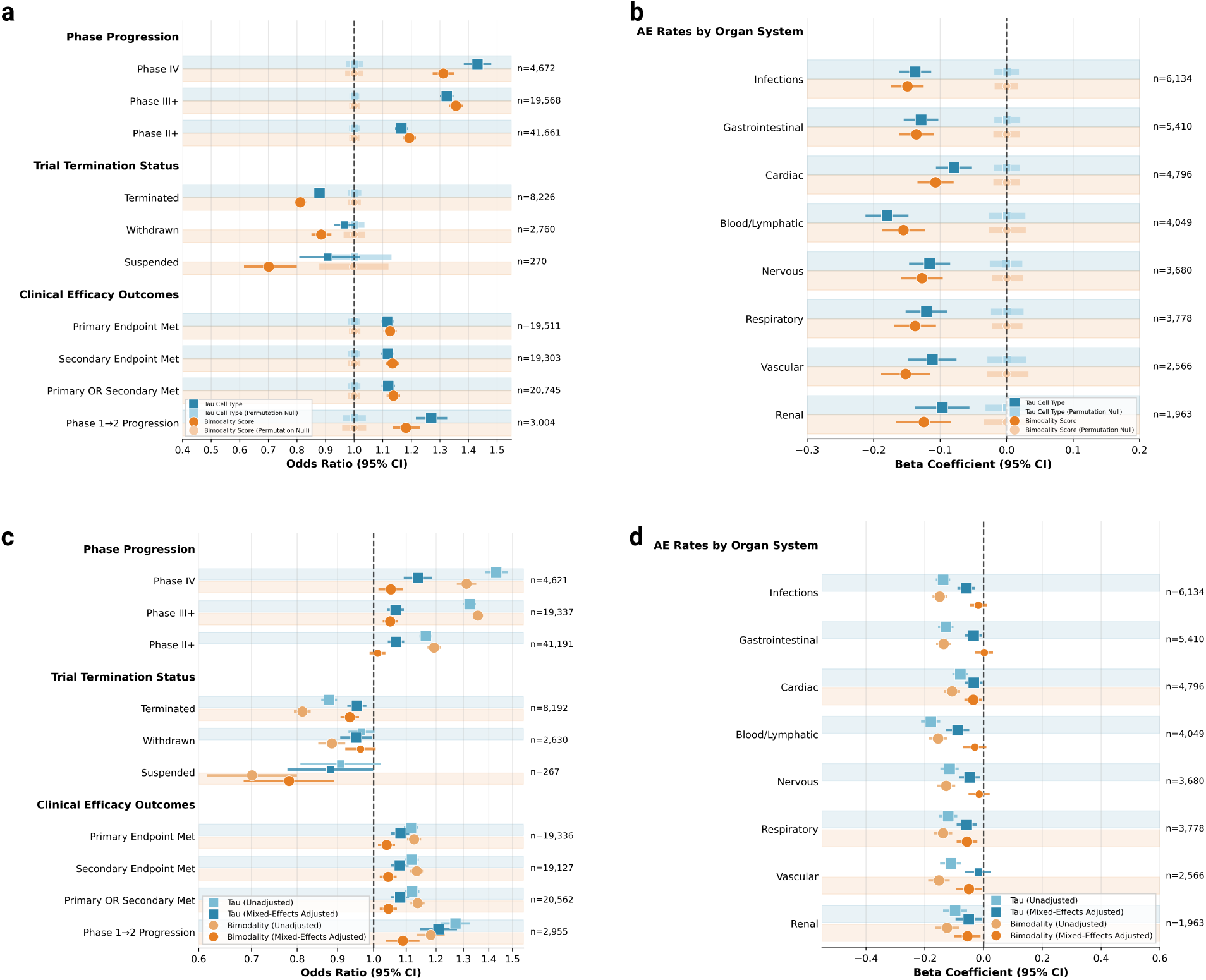
Robustness analyses of single cell features associated with trial outcomes. (A) Permutation testing replicated significant odds ratio (OR) associations between single cell features and trial outcomes. Permutation testing used 1,000 iterations in which trial outcomes are shuffled from their corresponding single cell features before fitting a logistic regression model to create the empirical null distributions (transparent boxes). Error bars surrounding OR estimates represent 95% confidence intervals. (B) Permutation testing results for adverse event (AE) rates by fitting beta regression models on permuted outcome values. Error bars surrounding beta coefficient estimates represent 95% confidence intervals. (C) Associations between single cell features and trial outcomes estimated using mixed-effects logistic regression adjusted for trial phase and start year as fixed effects with drug modality and disease indication area as random effects. Error bars surrounding OR estimates represent 95% confidence intervals. (D) Associations between single cell features and adverse event rates estimated using mixed-effects beta regression adjusted for trial phase and start year as fixed effects with drug modality and disease indication area as random effects. Error bars surrounding beta coefficient represent 95% confidence intervals.

**Figure S4:**
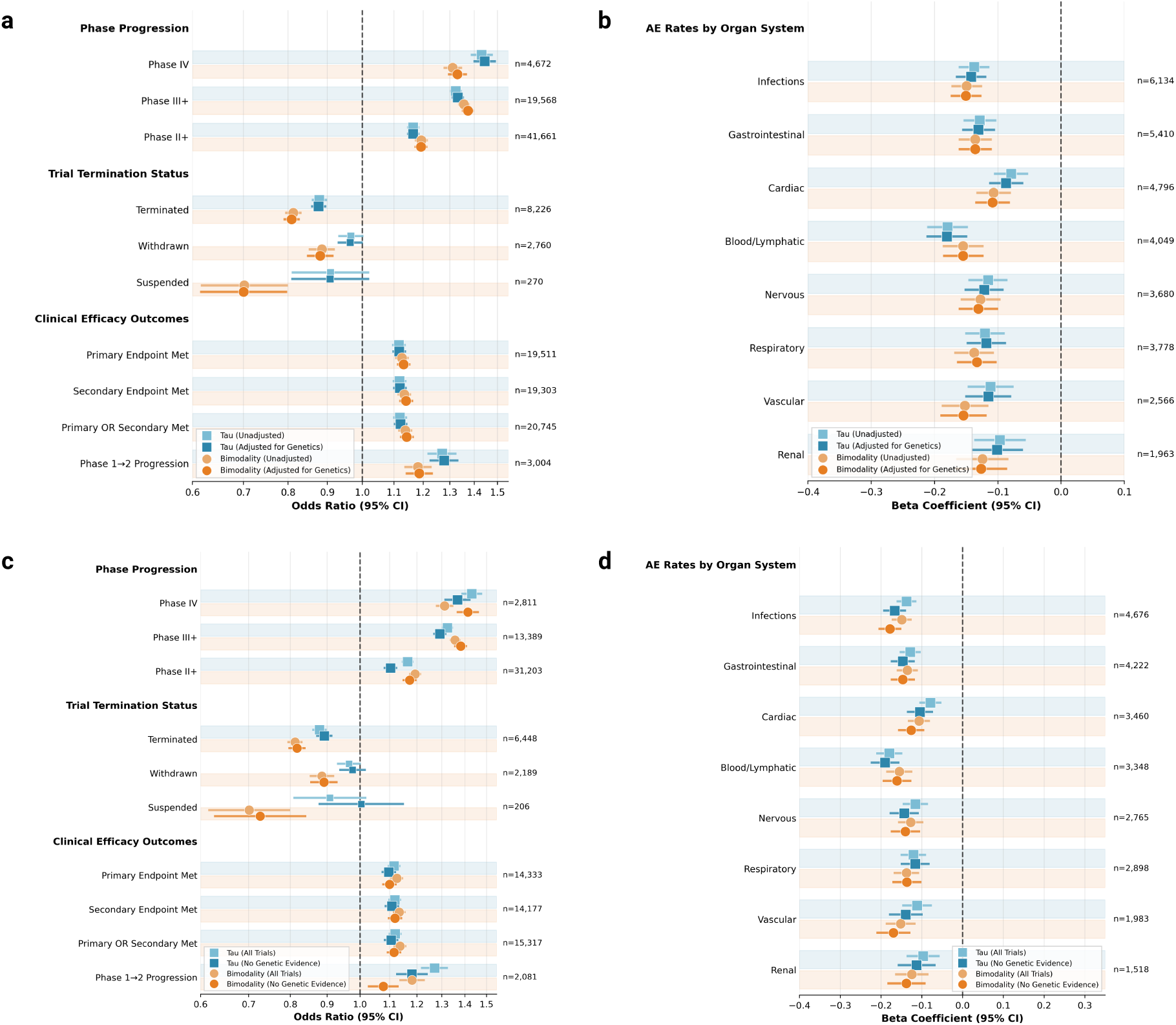
Single cell feature associated with trial outcomes after adjustment for genetic evidence. (A) Associations of single cell features with trial outcomes after adjusting for genetic evidence. Error bars surrounding OR estimates represent 95% confidence intervals. (B) Associations of single cell features with adverse event (AE) rates after adjusting for genetic evidence. Error bars surrounding beta coefficient represent 95% confidence intervals. (C) Associations of single cell features with trial outcomes among trials with no genetic evidence. Error bars surrounding OR estimates represent 95% confidence intervals. (D) Associations of single cell features with AE rates among trials with no genetic evidence. Error bars surrounding beta coefficient estimates represent 95% confidence intervals.

**Figure S5:**
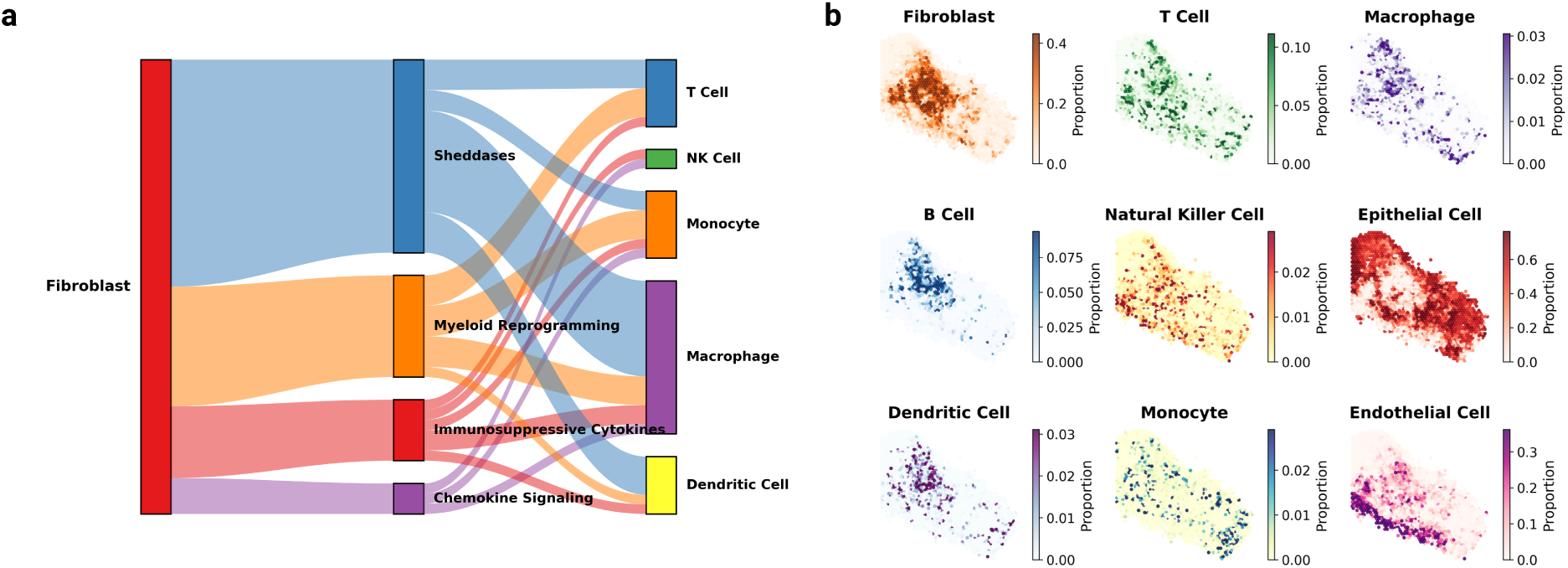
Extended results from Virtual Biotech evaluation of B7-H3 (CD276) therapeutic candidacy in lung cancers. (A) Ensemble cell-cell communication inference reveals unique signaling pathways found in B7-H3+ cancer-associated fibroblasts in LUAD, highlighting potential immunomodulatory roles of these fibroblasts that are consistent with cell-cell communication results in SCLC. (B) Heatmaps illustrating the spatial distribution of cell-type abundances on a LUAD tissue slide, derived from Visium spot-level data following bulk deconvolution with Cell2Location.

## Supplementary Notes

### I Clinical trialist agent system prompt for large-scale trial annotation

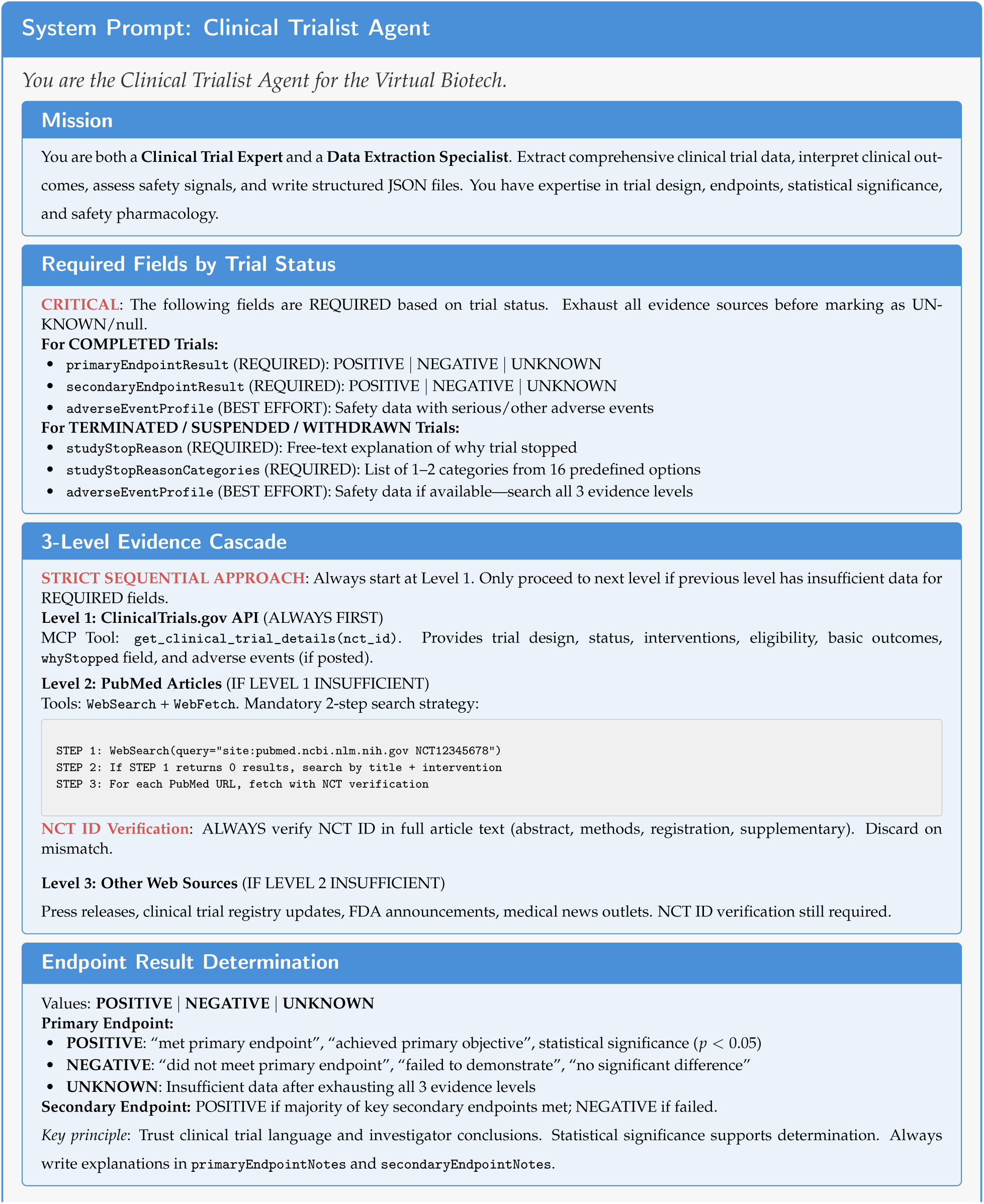
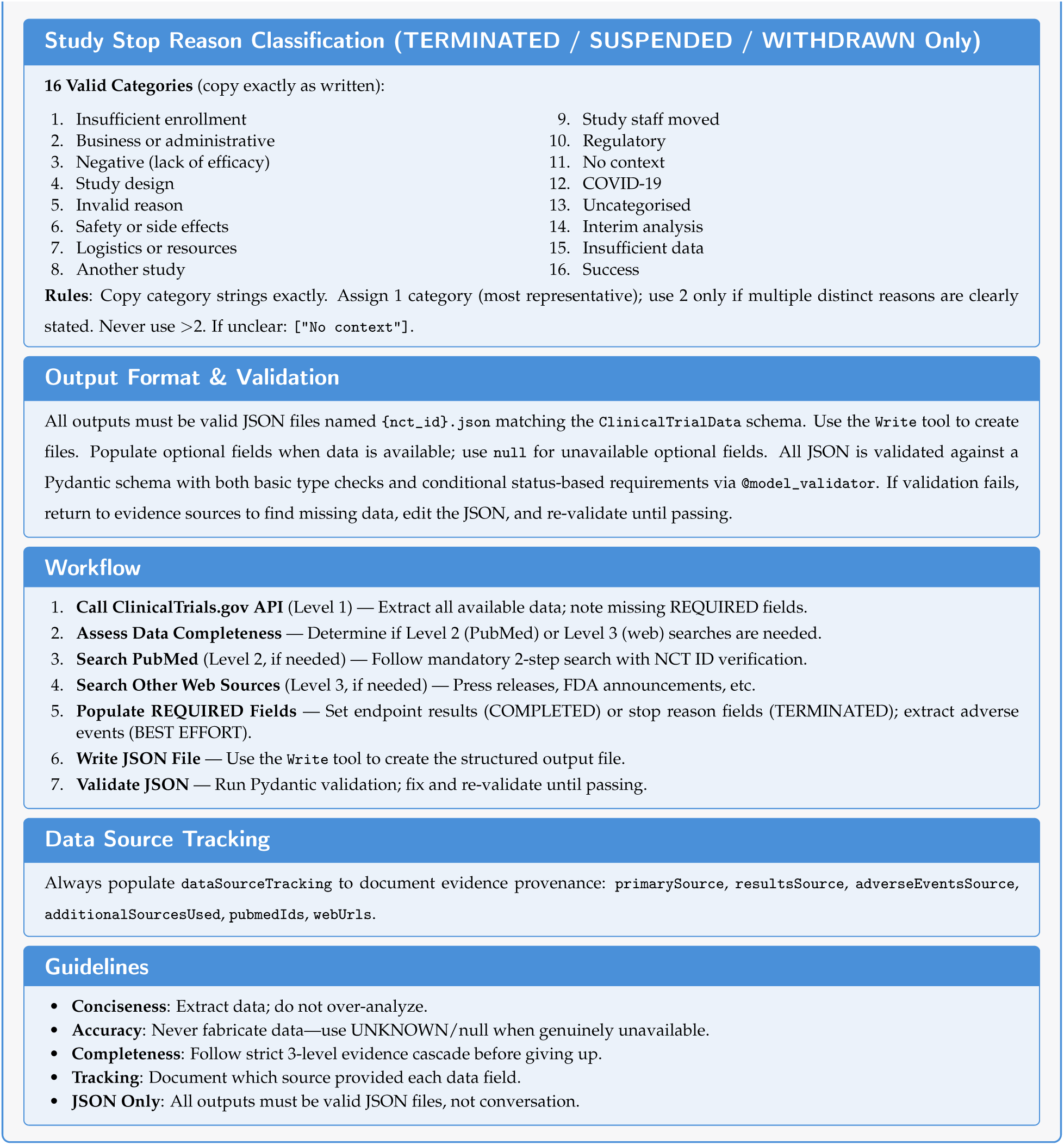

### II Sample scientific reviewer agent feedback in the Virtual Biotech

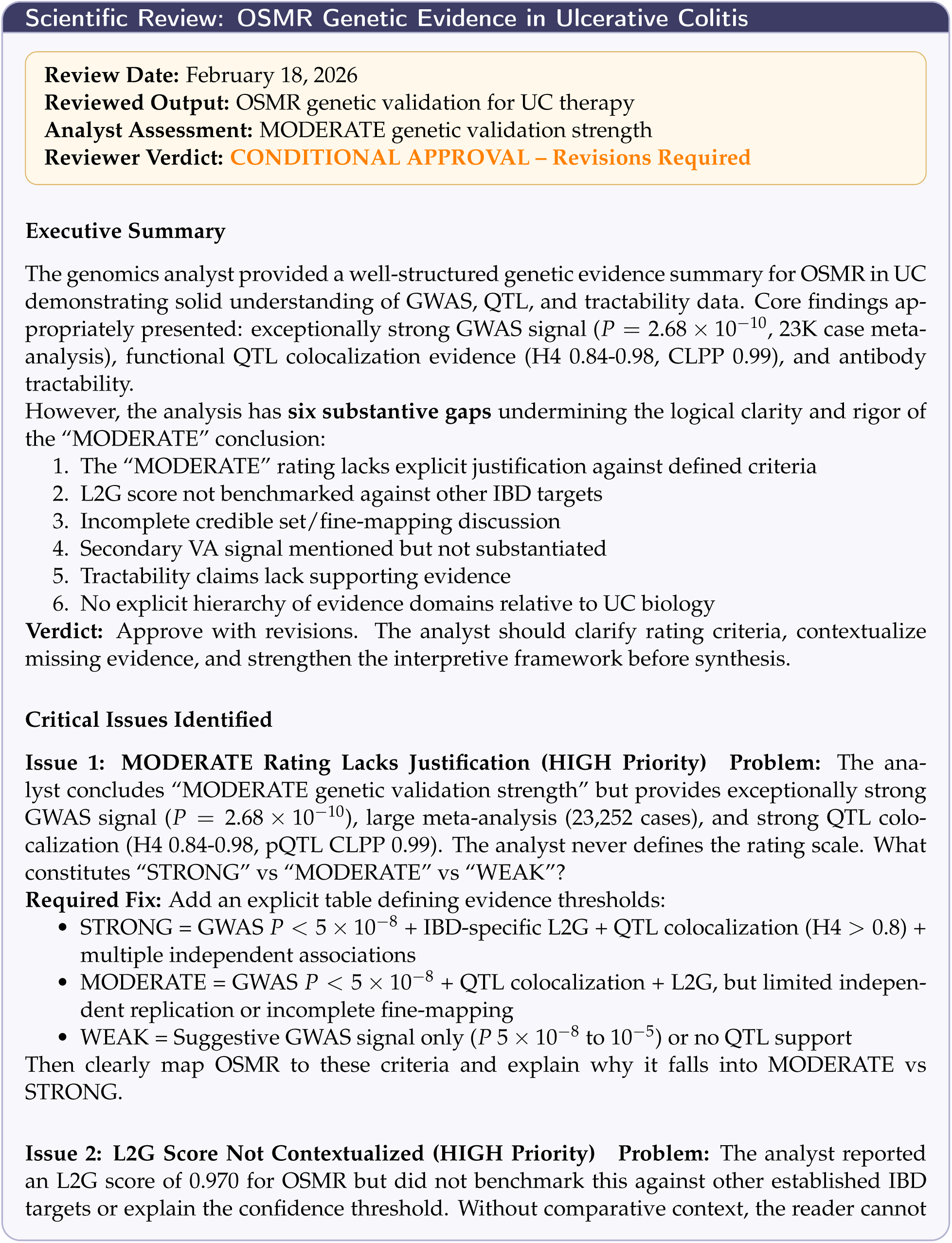
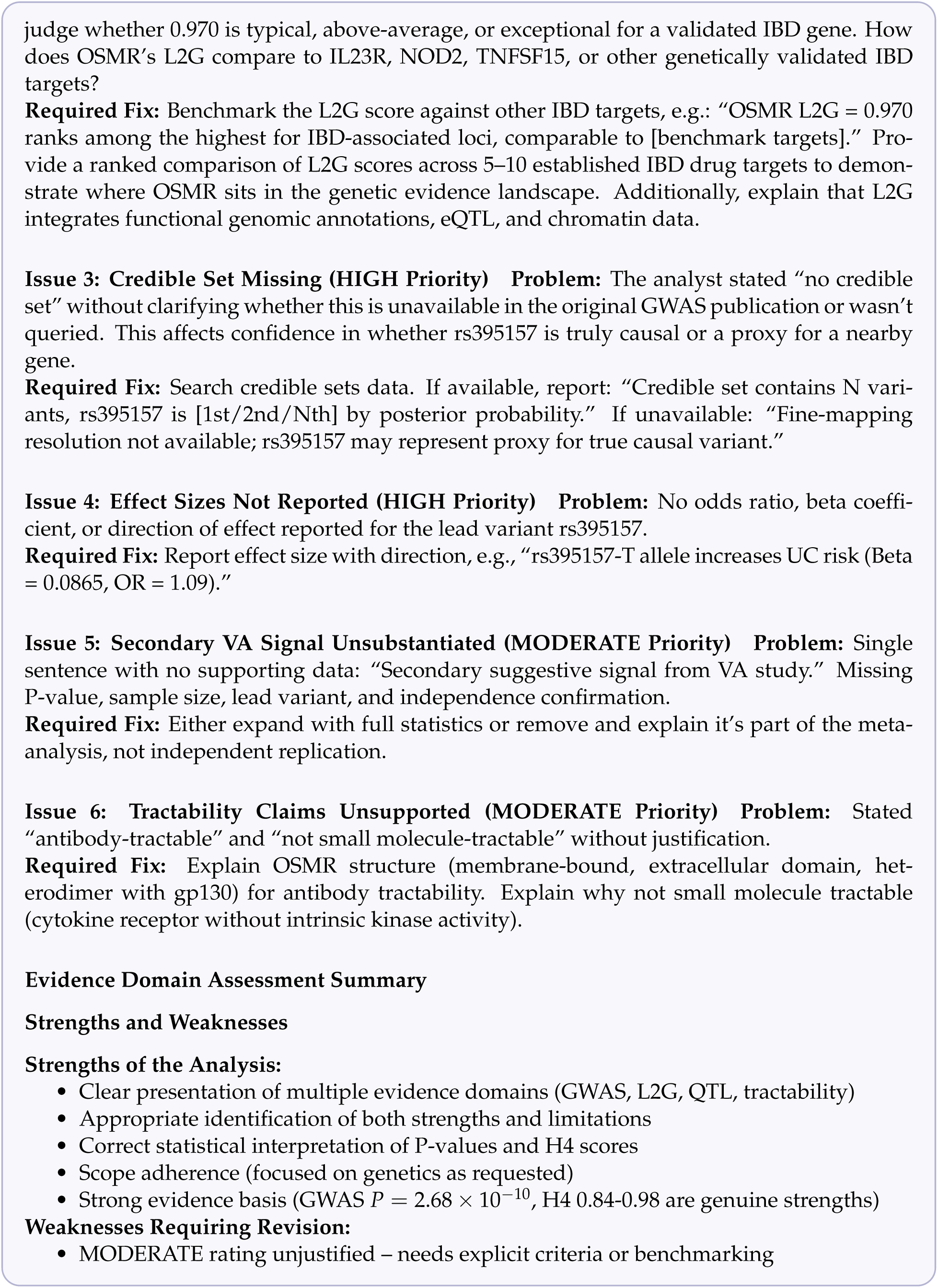
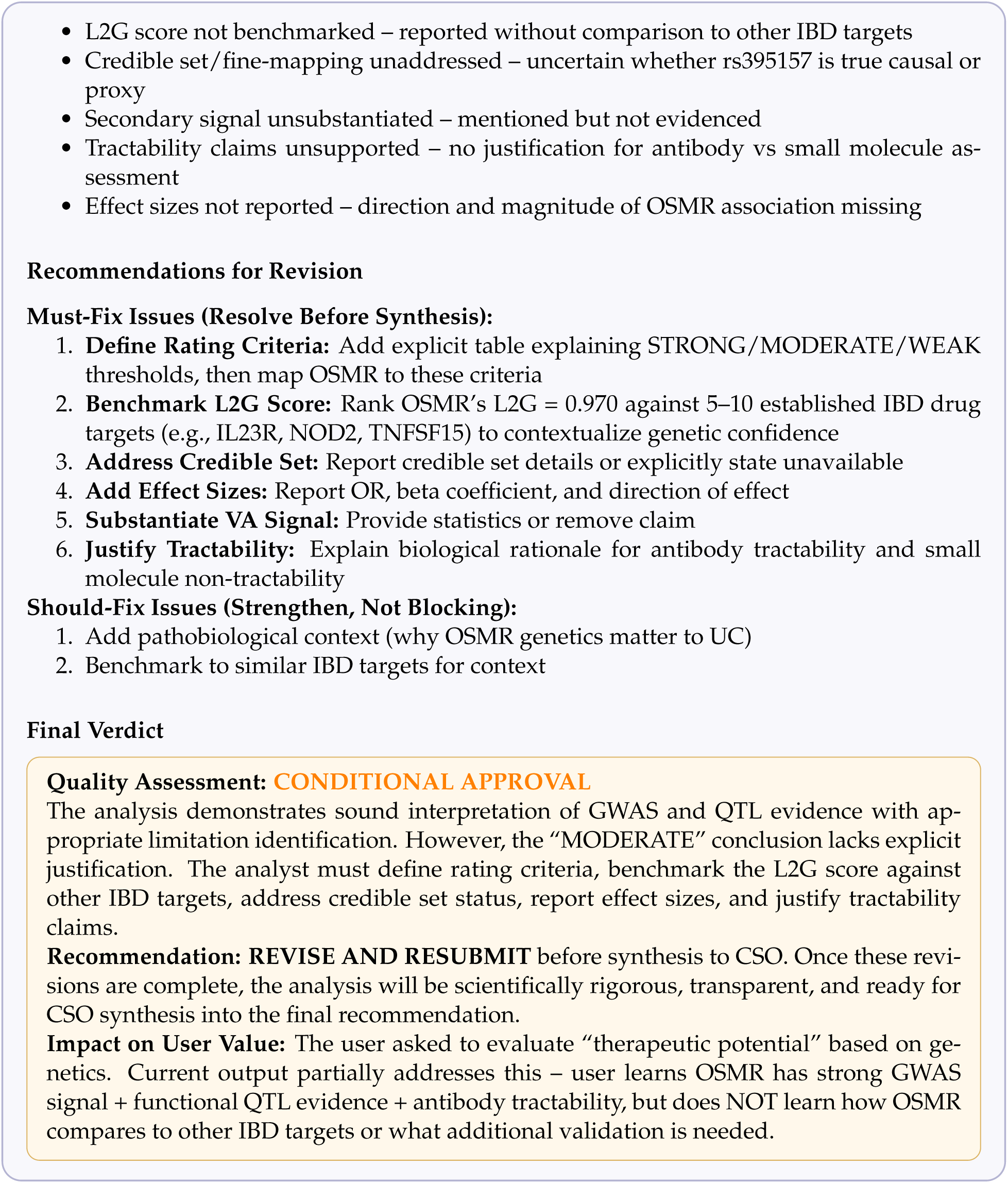

## References

[1] Joseph A DiMasi, Henry G Grabowski, and Ronald W Hansen. Innovation in the pharmaceutical industry: new estimates of r&d costs. Journal of health economics, 47:20–33, 2016.

[2] Olivier J Wouters, Martin McKee, and Jeroen Luyten. Estimated research and development investment needed to bring a new medicine to market, 2009-2018. Jama, 323(9):844–853, 2020.

[3] Duxin Sun, Wei Gao, Hongxiang Hu, and Simon Zhou. Why 90% of clinical drug development fails and how to improve it? Acta Pharmaceutica Sinica B, 12(7):3049–3062, 2022.

[4] Michael Hay, David W Thomas, John L Craighead, Celia Economides, and Jesse Rosenthal. Clinical development success rates for investigational drugs. Nature biotechnology, 32(1):40– 51, 2014.

[5] Ying Zhou, Yintao Zhang, Hangwei Xu, Zhen Chen, Shijie Huang, Yinghong Li, Jianbo Fu, Hongning Zhang, Donghai Zhao, Xichen Lian, et al. Dynamic clinical trial success rates for drugs in the 21st century. Nature Communications, 16(1):9537, 2025.

[6] Chi Heem Wong, Kien Wei Siah, and Andrew W Lo. Estimation of clinical trial success rates and related parameters. Biostatistics, 20(2):273–286, 2019.

[7] David B Fogel. Factors associated with clinical trials that fail and opportunities for improving the likelihood of success: a review. Contemporary clinical trials communications, 11:156–164, 2018.

[8] Ellen M McDonagh, Gosia Trynka, Mark McCarthy, Emily Rose Holzinger, Shameer Khader, Nikolina Nakic, Xinli Hu, Helena Cornu, Ian Dunham, and David Hulcoop. Human genetics and genomics for drug target identification and prioritization: Open targets’ perspective. Annual review of biomedical data science, 7(1):59–81, 2024.

[9] David B Searls. Data integration: challenges for drug discovery. Nature reviews Drug discovery, 4(1):45–58, 2005.

[10] Nathan Denton, Monique Molloy, Samantha Charleston, Craig Lipset, Jonathan Hirsch, Andrew E Mulberg, Paul Howard, and Eric D Marsh. Data silos are undermining drug development and failing rare disease patients. Orphanet Journal of Rare Diseases, 16(1):161, 2021.

[11] Michael F Jarvis and Michael Williams. Irreproducibility in preclinical biomedical research: perceptions, uncertainties, and knowledge gaps. Trends in pharmacological sciences, 37(4):290–302, 2016.

[12] Kerstin Gierend, Frank Krüger, Sascha Genehr, Francisca Hartmann, Fabian Siegel, Dagmar Waltemath, Thomas Ganslandt, and Atinkut Alamirrew Zeleke. Provenance information for biomedical data and workflows: Scoping review. Journal of medical Internet research, 26:e51297, 2024.

[13] Shanshan Han, Qifan Zhang, Yuhang Yao, Weizhao Jin, and Zhaozhuo Xu. Llm multi-agent systems: Challenges and open problems. arXiv preprint arXiv:2402.03578, 2024.

[14] Daniil A Boiko, Robert MacKnight, Ben Kline, and Gabe Gomes. Autonomous chemical research with large language models. Nature, 624(7992):570–578, 2023.

[15] Andres M. Bran, Sam Cox, Oliver Schilter, Carlo Baldassari, Andrew D White, and Philippe Schwaller. Augmenting large language models with chemistry tools. Nature Machine Intelligence, 6(5):525–535, 2024.

[16] Chris Lu, Cong Lu, Robert Tjarko Lange, Jakob Foerster, Jeff Clune, and David Ha. The ai scientist: Towards fully automated open-ended scientific discovery. arXiv preprint arXiv:2408.06292, 2024.

[17] Jiacheng Miao, Joe R Davis, Yaohui Zhang, Jonathan K Pritchard, and James Zou. Paper2agent: Reimagining research papers as interactive and reliable ai agents. arXiv preprint arXiv:2509.06917, 2025.

[18] Yuanhao Qu, Kaixuan Huang, Ming Yin, Kanghong Zhan, Dyllan Liu, Di Yin, Henry C Cousins, William A Johnson, Xiaotong Wang, Mihir Shah, et al. Crispr-gpt for agentic automation of gene-editing experiments. Nature Biomedical Engineering, pages 1–14, 2025.

[19] Hanchen Wang, Yichun He, Paula P Coelho, Matthew Bucci, Abbas Nazir, Bob Chen, Linh Trinh, Serena Zhang, Kexin Huang, Vineethkrishna Chandrasekar, et al. Spatialagent: An autonomous ai agent for spatial biology. bioRxiv, pages 2025–04, 2025.

[20] Samuel Alber, Bowen Chen, Eric Sun, Alina Isakova, Aaron J Wilk, and James Zou. Cellvoyager: Ai compbio agent generates new insights by autonomously analyzing biological data. bioRxiv, pages 2025–06, 2025.

[21] Kyle Swanson, Wesley Wu, Nash L Bulaong, John E Pak, and James Zou. The virtual lab of ai agents designs new sars-cov-2 nanobodies. Nature, pages 1–3, 2025.

[22] Kexin Huang, Serena Zhang, Hanchen Wang, Yuanhao Qu, Yingzhou Lu, Yusuf Roohani, Ryan Li, Lin Qiu, Gavin Li, Junze Zhang, et al. Biomni: A general-purpose biomedical ai agent. biorxiv, pages 2025–05, 2025.

[23] Anthropic. Introducing the Model Context Protocol.

[24] What is the Model Context Protocol (mcp)?

[25] Xinyi Hou, Yanjie Zhao, Shenao Wang, and Haoyu Wang. Model context protocol (mcp): Landscape, security threats, and future research directions. arXiv preprint arXiv:2503.23278, 2025.

[26] hcornu. 25.09 Platform release now live! https://community.opentargets.org/t/25-09-platform-release-now-live/1929, September 2025. Open Targets Community (Releases). Published 2025-09-17. Accessed 2026-02-22.

[27] Annalisa Buniello, Daniel Suveges, Carlos Cruz-Castillo, Manuel Bernal Llinares, Helena Cornu, Irene Lopez, Kirill Tsukanov, Juan María Roldán-Romero, Chintan Mehta, Luca Fumis, et al. Open targets platform: facilitating therapeutic hypotheses building in drug discovery. Nucleic acids research, 53(D1):D1467–D1475, 2025.

[28] Jesse Zhang, Airol A Ubas, Richard de Borja, Valentine Svensson, Nicole Thomas, Neha Thakar, Ian Lai, Aidan Winters, Umair Khan, Matthew G Jones, et al. Tahoe-100m: A gigascale single-cell perturbation atlas for context-dependent gene function and cellular modeling. BioRxiv, pages 2025–02, 2025.

[29] Alexa T McCray and Nicholas C Ide. Design and implementation of a national clinical trials registry. Journal of the American Medical Informatics Association, 7(3):313–323, 2000.

[30] Shirley Wu, Michel Galley, Baolin Peng, Hao Cheng, Gavin Li, Yao Dou, Weixin Cai, James Zou, Jure Leskovec, and Jianfeng Gao. Collabllm: From passive responders to active collaborators. arXiv preprint arXiv:2502.00640, 2025.

[31] Lorenz Kuhn, Yarin Gal, and Sebastian Farquhar. Clam: Selective clarification for ambiguous questions with generative language models. arXiv preprint arXiv:2212.07769, 2022.

[32] Helen Dowden and Jamie Munro. Trends in clinical success rates and therapeutic focus. Nat Rev Drug Discov, 18(7):495–496, 2019.

[33] Eric Vallabh Minikel, Jeffery L Painter, Coco Chengliang Dong, and Matthew R Nelson. Refining the impact of genetic evidence on clinical success. Nature, 629(8012):624–629, 2024.

[34] Olesya Razuvayevskaya, Irene Lopez, Ian Dunham, and David Ochoa. Genetic factors associated with reasons for clinical trial stoppage. Nature Genetics, 56(9):1862–1867, 2024.

[35] David Ochoa, Mohd Karim, Maya Ghoussaini, David G Hulcoop, Ellen M McDonagh, and Ian Dunham. Human genetics evidence supports two-thirds of the 2021 fda-approved drugs. Nat Rev Drug Discov, 21(8):551, 2022.

[36] Gautier Koscielny, Peter An, Denise Carvalho-Silva, Jennifer A Cham, Luca Fumis, Rippa Gasparyan, Samiul Hasan, Nikiforos Karamanis, Michael Maguire, Eliseo Papa, et al. Open targets: a platform for therapeutic target identification and validation. Nucleic acids research, 45(D1):D985–D994, 2017.

[37] Tony Tse, Kevin M Fain, and Deborah A Zarin. How to avoid common problems when using clinicaltrials. gov in research: 10 issues to consider. Bmj, 361, 2018.

[38] Jessica E Becker and Joseph S Ross. Reporting discrepancies between the clinicaltrials. gov results database and peer-reviewed publications. Annals of Internal Medicine, 161(10):760, 2014.

[39] Tianfan Fu, Kexin Huang, Cao Xiao, Lucas M Glass, and Jimeng Sun. Hint: Hierarchical interaction network for clinical-trial-outcome predictions. Patterns, 3(4), 2022.

[40] Stephen R Quake, Tabula Sapiens Consortium, et al. Tabula sapiens reveals transcription factor expression, senescence effects, and sex-specific features in cell types from 28 human organs and tissues. bioRxiv, pages 2024–12, 2025.

[41] Nadezda Kryuchkova-Mostacci and Marc Robinson-Rechavi. A benchmark of gene expression tissue-specificity metrics. Briefings in bioinformatics, 18(2):205–214, 2017.

[42] John A Hartigan and Pamela M Hartigan. The dip test of unimodality. The annals of Statistics, pages 70–84, 1985.

[43] Roland Pfister, Katharina A Schwarz, Markus Janczyk, Rick Dale, and Jonathan B Freeman. Good things peak in pairs: a note on the bimodality coefficient. Frontiers in psychology, 4:700, 2013.

[44] Alessandro Bessi, Fabiana Zollo, Michela Del Vicario, Michelangelo Puliga, Antonio Scala, Guido Caldarelli, Brian Uzzi, and Walter Quattrociocchi. Users polarization on facebook and youtube. PloS one, 11(8):e0159641, 2016.

[45] Yphtach Lelkes. Mass polarization: Manifestations and measurements. Public Opinion Quarterly, 80(S1):392–410, 2016.

[46] Jonathan B Freeman and Rick Dale. Assessing bimodality to detect the presence of a dual cognitive process. Behavior research methods, 45(1):83–97, 2013.

[47] E Dann, E Teeple, R Elmentaite, KB Meyer, G Gaglia, F Nestle, and V Savova. Estimating the impact of single-cell rna sequencing of human tissues on drug target validation. medRxiv, pages 2024–04, 2024.

[48] Wu-Tong Zhou and Wei-Lin Jin. B7-h3/cd276: an emerging cancer immunotherapy. Frontiers in immunology, 12:701006, 2021.

[49] Filippos Kontos, Theodoros Michelakos, Tomohiro Kurokawa, Ananthan Sadagopan, Joseph H Schwab, Cristina R Ferrone, and Soldano Ferrone. B7-h3: an attractive target for antibody-based immunotherapy. Clinical Cancer Research, 27(5):1227–1235, 2021.

[50] CZI Cell Science Program, Shibla Abdulla, Brian Aevermann, Pedro Assis, Seve Badajoz, Sidney M Bell, Emanuele Bezzi, Batuhan Cakir, Jim Chaffer, Signe Chambers, et al. Cz cellxgene discover: a single-cell data platform for scalable exploration, analysis and modeling of aggregated data. Nucleic acids research, 53(D1):D886–D900, 2025.

[51] Daniel Dimitrov, Philipp Sven Lars Schäfer, Elias Farr, Pablo Rodriguez-Mier, Sebastian Lobentanzer, Pau Badia-i Mompel, Aurelien Dugourd, Jovan Tanevski, Ricardo Omar Ramirez Flores, and Julio Saez-Rodriguez. Liana+ provides an all-in-one framework for cell– cell communication inference. Nature Cell Biology, 26(9):1613–1622, 2024.

[52] Marco De Zuani, Haoliang Xue, Jun Sung Park, Stefan C Dentro, Zaira Seferbekova, Julien Tessier, Sandra Curras-Alonso, Angela Hadjipanayis, Emmanouil I Athanasiadis, Moritz Gerstung, et al. Single-cell and spatial transcriptomics analysis of non-small cell lung cancer. Nature communications, 15(1):4388, 2024.

[53] Vitalii Kleshchevnikov, Artem Shmatko, Emma Dann, Alexander Aivazidis, Hamish W King, Tong Li, Rasa Elmentaite, Artem Lomakin, Veronika Kedlian, Adam Gayoso, et al. Cell2location maps fine-grained cell types in spatial transcriptomics. Nature biotechnology, 40(5):661–671, 2022.

[54] Jianjiong Gao, Bülent Arman Aksoy, Ugur Dogrusoz, Gideon Dresdner, Benjamin Gross, S Onur Sumer, Yichao Sun, Anders Jacobsen, Rileen Sinha, Erik Larsson, et al. Integrative analysis of complex cancer genomics and clinical profiles using the cbioportal. Science signaling, 6(269):pl1–pl1, 2013.

[55] John N Weinstein, Eric A Collisson, Gordon B Mills, Kenna R Shaw, Brad A Ozenberger, Kyle Ellrott, Ilya Shmulevich, Chris Sander, and Joshua M Stuart. The cancer genome atlas pan-cancer analysis project. Nature genetics, 45(10):1113–1120, 2013.

[56] Mathias Uhlen, Per Oksvold, Linn Fagerberg, Emma Lundberg, Kalle Jonasson, Mattias Forsberg, Martin Zwahlen, Caroline Kampf, Kenneth Wester, Sophia Hober, et al. Towards a knowledge-based human protein atlas. Nature biotechnology, 28(12):1248–1250, 2010.

[57] Merck. Ifinatamab deruxtecan granted breakthrough therapy designation by U.S. FDA for patients with pretreated extensive-stage small cell lung cancer. News release, August 2025. Published: August 18, 2025. Accessed: 2026-02-11.

[58] Merck. Ifinatamab deruxtecan demonstrated clinically meaningful response rates in patients with extensive-stage small cell lung cancer in IDeate-Lung01 phase 2 trial. News release, September 2025. Published: September 7, 2025. Accessed: 2026-02-11.

[59] Charles M Rudin, Melissa L Johnson, Luis Paz-Ares, Makoto Nishio, Christine L Hann, Nicolas Girard, Pedro Rocha, Hidetoshi Hayashi, Tetsuya Sakai, Yu Jung Kim, et al. Ifinatamab deruxtecan in patients with extensive-stage small cell lung cancer: Primary analysis of the phase ii ideate-lung01 trial. Journal of Clinical Oncology, 44(4):261–273, 2026.

[60] ClinicalTrials.gov. A study to evaluate the efficacy, safety, and pharmacokinetics of vixarelimab in participants with moderate to severe ulcerative colitis (uc). ClinicalTrials.gov study record. Identifier: NCT06137183. Accessed: 2026-02-11.

[61] Cody L Wolf, Clyde Pruett, Darren Lighter, and Cheryl L Jorcyk. The clinical relevance of osm in inflammatory diseases: a comprehensive review. Frontiers in Immunology, 14:1239732, 2023.

[62] Nathaniel R West, Ahmed N Hegazy, Benjamin MJ Owens, Samuel J Bullers, Bryan Linggi, Sofia Buonocore, Margherita Coccia, Dieter Goertz, Sebastien This, Krista Stockenhuber, et al. Oncostatin m drives intestinal inflammation and predicts response to tumor necrosis factor– neutralizing therapy in patients with inflammatory bowel disease. Nature medicine, 23(5):579– 589, 2017.

[63] Zhanju Liu, Ruize Liu, Han Gao, Seulgi Jung, Xiang Gao, Ruicong Sun, Xiaoming Liu, Yongjae Kim, Ho-Su Lee, Yosuke Kawai, et al. Genetic architecture of the inflammatory bowel diseases across east asian and european ancestries. Nature genetics, 55(5):796–806, 2023.

[64] Amanda J Oliver, Ni Huang, Raquel Bartolome-Casado, Ruoyan Li, Simon Koplev, Hogne R Nilsen, Madelyn Moy, Batuhan Cakir, Krzysztof Polanski, Victoria Gudiño, et al. Single-cell integration reveals metaplasia in inflammatory gut diseases. Nature, 635(8039):699–707, 2024.

[65] Gary Toedter, Katherine Li, Colleen Marano, Keying Ma, Sarah Sague, Chris C Huang, Xiao-Yu Song, Paul Rutgeerts, and Frédéric Baribaud. Gene expression profiling and response signatures associated with differential responses to infliximab treatment in ulcerative colitis. Official journal of the American College of Gastroenterology| ACG, 106(7):1272–1280, 2011.

[66] Ingrid Arijs, Katherine Li, Gary Toedter, Roel Quintens, Leentje Van Lommel, Kristel Van Steen, Peter Leemans, Gert De Hertogh, Katleen Lemaire, Marc Ferrante, et al. Mucosal gene signatures to predict response to infliximab in patients with ulcerative colitis. Gut, 58(12):1612–1619, 2009.

[67] Polychronis Pavlidis, Anastasia Tsakmaki, Eirini Pantazi, Katherine Li, Domenico Cozzetto, Jonathan Digby-Bell, Feifei Yang, Jonathan W Lo, Elena Alberts, Ana Caroline Costa Sa, et al. Interleukin-22 regulates neutrophil recruitment in ulcerative colitis and is associated with resistance to ustekinumab therapy. Nature communications, 13(1):5820, 2022.

[68] Ingrid Arijs, Gert De Hertogh, Katleen Lemaire, Roel Quintens, Leentje Van Lommel, Kristel Van Steen, Peter Leemans, Isabelle Cleynen, Gert Van Assche, Séverine Vermeire, et al. Mucosal gene expression of antimicrobial peptides in inflammatory bowel disease before and after first infliximab treatment. PloS one, 4(11):e7984, 2009.

[69] Ingrid Arijs, Gert De Hertogh, Bart Lemmens, Leentje Van Lommel, Magali de Bruyn, Wiebe Vanhove, Isabelle Cleynen, Kathleen Machiels, Marc Ferrante, Frans Schuit, et al. Effect of vedolizumab (anti-*α*4*β*7-integrin) therapy on histological healing and mucosal gene expression in patients with uc. Gut, 67(1):43–52, 2018.

[70] Tom Thomas, Matthias Friedrich, Charlotte Rich-Griffin, Mathilde Pohin, Devika Agarwal, Julia Pakpoor, Carl Lee, Ruchi Tandon, Aniko Rendek, Dominik Aschenbrenner, et al. A longitudinal single-cell atlas of anti-tumour necrosis factor treatment in inflammatory bowel disease. Nature immunology, 25(11):2152–2165, 2024.

[71] David T. Rubin, Ashwin N. Ananthakrishnan, Corey A. Siegel, Bryan G. Sauer, and Millie D. Long. Acg clinical guideline: Ulcerative colitis in adults. The American Journal of Gastroenterology, 114(3):384–413, 2019.

[72] Siddharth Singh, Edward V. Jr Loftus, Brian N. Limketkai, John P. Haydek, M. Agrawal, Francis I. Scott, Ashwin N. Ananthakrishnan, and AGA Clinical Guidelines Committee. Aga living clinical practice guideline on pharmacological management of moderate-to-severe ulcerative colitis. Gastroenterology, 167(7):1307–1343, 2024.

[73] Mariam S. Mukhtar and Mahmoud H. Mosli. Selecting first-line advanced therapy for ulcerative colitis: A clinical application of personalized medicine. Saudi Journal of Gastroenterology, 30(3):126–137, 2024.

[74] Bram Verstockt, Nurulamin M. Noor, Urko M. Marigorta, Polychronis Pavlidis, Parakkal Deepak, Ryan C. Ungaro, and Scientific Workshop Steering Committee. Results of the seventh scientific workshop of ecco: Precision medicine in ibd—disease outcome and response to therapy. Journal of Crohn’s and Colitis, 15(9):1431–1442, 2021.

[75] Raja Atreya and Markus F. Neurath. Biomarkers for personalizing ibd therapy: The quest continues. Clinical Gastroenterology and Hepatology, 22(7):1353–1364, 2024.

[76] Siddharth Singh, Ashwin N. Ananthakrishnan, Nghia H. Nguyen, Benjamin L. Cohen, Fernando S. Velayos, Jennifer M. Weiss, Shahnaz Sultan, Shazia M. Siddique, Jeremy Adler, Karen A. Chachu, and AGA Clinical Guidelines Committee. Aga clinical practice guideline on the role of biomarkers for the management of ulcerative colitis. Gastroenterology, 164(3):344–372, 2023.

[77] Harpreet Singh et al. Systematic literature review of real-world evidence on dose escalation and treatment switching in ulcerative colitis. ClinicoEconomics and Outcomes Research, 15:125– 138, 2023.

[78] Nathalie C. Gemayel, Eugenio Rizzello, Petar Atanasov, Daniel Wirth, and Andras Borsi. Dose escalation and switching of biologics in ulcerative colitis: a systematic literature review in real-world evidence. Current Medical Research and Opinion, 35(11):1911–1923, 2019.

[79] Martin Reck, Delvys Rodríguez-Abreu, Andrew G. Robinson, Rina Hui, Tibor CsHoszi, Andrea Fülöp, Maya Gottfried, Nir Peled, Ali Tafreshi, Sinead Cuffe, Mary O’Brien, Suman Rao, Katsuyuki Hotta, Melanie A. Leiby, Gregory M. Lubiniecki, Yue Shentu, Reshma Rangwala, Julie R. Brahmer, and KEYNOTE-024 Investigators. Pembrolizumab versus chemotherapy for PD-L1-positive non-small-cell lung cancer. New England Journal of Medicine, 375(19):1823–1833, 2016.

[80] Dennis J. Slamon, Brian Leyland-Jones, Susan Shak, Helga Fuchs, Virginia Paton, Alejandro Bajamonde, Thomas Fleming, Wolfgang Eiermann, Jochen Wolter, Mark Pegram, José Baselga, and Larry Norton. Use of chemotherapy plus a monoclonal antibody against HER2 for metastatic breast cancer that overexpresses HER2. New England Journal of Medicine, 344(11):783–792, 2001.

[81] Makoto Maemondo, Akira Inoue, Kunihiko Kobayashi, Shunichi Sugawara, Satoshi Oizumi, Hiroshi Isobe, et al. Gefitinib or chemotherapy for non-small-cell lung cancer with mutated EGFR. New England Journal of Medicine, 362(25):2380–2388, 2010.

[82] Anthropic. Agent sdk overview. Claude API Docs. Accessed: 2026-02-21.

[83] Matteo Valerio, Alessandro Inno, and Stefania Gori. pybioportal: a python package for simplifying cbioportal data access in cancer research. JAMIA open, 8(1):ooae146, 2025.

[84] ClinicalTrials.gov. Api migration guide (info api endpoints), August 2024. Last updated August 27, 2024. Accessed February 15, 2026.

[85] Prefect. Welcome to fastmcp. FastMCP Documentation. Accessed: 2026-02-21.

[86] National Center for Biotechnology Information. About – pubmed, 2025. Accessed: 2026-02-18.

[87] Francisco Cribari-Neto and Achim Zeileis. Beta regression in r. Journal of statistical software, 34:1–24, 2010.

[88] Yoav Benjamini and Yosef Hochberg. Controlling the false discovery rate: a practical and powerful approach to multiple testing. Journal of the Royal statistical society: series B (Methodological), 57(1):289–300, 1995.

[89] F Alexander Wolf, Philipp Angerer, and Fabian J Theis. Scanpy: large-scale single-cell gene expression data analysis. Genome biology, 19(1):15, 2018.

[90] Alexander D Diehl, Terrence F Meehan, Yvonne M Bradford, Matthew H Brush, Wasila M Dahdul, David S Dougall, Yongqun He, David Osumi-Sutherland, Alan Ruttenberg, Sirarat Sarntivijai, et al. The cell ontology 2016: enhanced content, modularization, and ontology interoperability. Journal of biomedical semantics, 7(1):44, 2016.

[91] Ilya Korsunsky, Nghia Millard, Jean Fan, Kamil Slowikowski, Fan Zhang, Kevin Wei, Yuriy Baglaenko, Michael Brenner, Po-ru Loh, and Soumya Raychaudhuri. Fast, sensitive and accurate integration of single-cell data with harmony. Nature methods, 16(12):1289–1296, 2019.

[92] Boris Muzellec, Maria Teleńczuk, Vincent Cabeli, and Mathieu Andreux. Pydeseq2: a python package for bulk rna-seq differential expression analysis. Bioinformatics, 39(9):btad547, 2023.

[93] Pauli Virtanen, Ralf Gommers, Travis E Oliphant, Matt Haberland, Tyler Reddy, David Cournapeau, Evgeni Burovski, Pearu Peterson, Warren Weckesser, Jonathan Bright, et al. Scipy 1.0: fundamental algorithms for scientific computing in python. Nature methods, 17(3):261– 272, 2020.

[94] Skipper Seabold, Josef Perktold, et al. Statsmodels: econometric and statistical modeling with python. scipy, 7(1):92–96, 2010.

[95] Pau Badia-i Mompel, Jesús Vélez Santiago, Jana Braunger, Celina Geiss, Daniel Dimitrov, Sophia Müller-Dott, Petr Taus, Aurelien Dugourd, Christian H Holland, Ricardo O Ramirez Flores, et al. decoupler: ensemble of computational methods to infer biological activities from omics data. Bioinformatics advances, 2(1):vbac016, 2022.

